# A Naïve RNA Sampling Core Enables Adaptive piRNA Specificity Against Transposable Elements

**DOI:** 10.64898/2026.02.07.704324

**Authors:** Dominik Handler, W. Y. Wylan Wong, Alexander Tsarev, Rippei Hayashi, Julius Brennecke

## Abstract

How piRNA-mediated genome defense achieves specificity against transposons while sampling a complex transcriptome has remained unresolved. Here we show that piRNA biogenesis operates through pervasive, non-specific sampling of cytoplasmic RNAs, with specificity imposed by tissue-specific molecular modules that exploit intrinsic vulnerabilities of transposons. In Drosophila somatic cells, the specificity factor Yb steers basal processing towards uridine-rich RNAs—automatically capturing antisense retrotransposon transcripts due to their intrinsically adenosine-biased genomes. In germline cells lacking Yb, basal sampling generates naïve piRNAs loaded into catalytically active Argonaute proteins, which trigger autocatalytic ping-pong amplification upon encountering complementary targets. In both contexts, transposon mobility facilitates the production of antisense RNAs that enable either biased processing or amplification. Thus, piRNA clusters, long associated with pathway specificity, act as sources of transposon antisense sequences, while specificity arises from layering distinct molecular mechanisms onto a shared foundation of indiscriminate transcript sampling, enabling robust and adaptable genome defense without predefined templates.

## INTRODUCTION

The PIWI-interacting RNA (piRNA) pathway is a conserved genome defense system that protects animal germlines from transposable elements through sequence-specific targeting by small RNAs loaded into PIWI-clade Argonaute proteins (Siomi et al., 2011, Ozata et al., 2018, Czech et al., 2018). A central unresolved question is how this pathway achieves specificity—preferentially generating piRNAs targeting transposon sequences—while remaining sufficiently flexible to respond to newly arising or horizontally acquired elements.

Unlike double-stranded RNA-triggered defense systems (Hannon, 2002, Ghildiyal and Zamore, 2009), piRNAs derive from single-stranded precursors (Vagin et al., 2006, Saito et al., 2006, Aravin et al., 2006, Girard et al., 2006, Lau et al., 2006, Brennecke et al., 2007, Gunawardane et al., 2007). This poses a conceptual challenge: antisense transposon transcripts must be distinguished from the vast excess of cellular mRNAs and selectively routed into piRNA biogenesis. Because transposons rarely encode their own antisense RNAs, such transcripts typically arise only when transposon sequences are embedded in host transcription units. The most prominent sources are piRNA clusters—genomic regions densely populated by transposon fragments that account for the majority of piRNA production and are widely viewed as heritable records of past transposon invasions (Aravin et al., 2003, Brennecke et al., 2007, Aravin et al., 2007).

These observations gave rise to the prevailing cluster-centric “trap-model”, which posits that pathway specificity is achieved by selectively recognizing cluster-derived transcripts through specialized sequence features, structural properties, or molecular marks acquired during nuclear processing (Senti and Brennecke, 2010, Zhang et al., 2014, Yu et al., 2019, Ishizu et al., 2015, Homolka et al., 2015, Takase et al., 2022). By jumping into a cluster, a transposon would thereby automatically expose its sequence to the piRNA machinery. However, despite extensive efforts, no universal features uniquely defining cluster transcripts have emerged. However, despite extensive efforts, no universal features uniquely defining cluster transcripts have emerged. Moreover, antisense transposon insertions in actively transcribed host genes outside canonical clusters can trigger robust piRNA production and effective silencing (Konstantinidou et al., 2024, Yu et al., 2025, Rafanel et al., 2025), indicating that cluster identity alone cannot account for specificity.

One established specificity mechanism is the ping-pong amplification cycle, in which piRNA-guided cleavage (slicing) of complementary targets generates new piRNAs in a self-reinforcing loop (Brennecke et al., 2007, Gunawardane et al., 2007, Aravin et al., 2007). Beyond ping-pong, piRNA-guided cleavage also initiates phased, 5′-to-3′ production of piRNAs along the target RNA (Mohn et al., 2015, Han et al., 2015, Homolka et al., 2015). However, slicing-induced piRNA biogenesis cannot explain how specificity arises in systems lacking catalytically competent PIWI proteins(Lau et al., 2009, Robine et al., 2009, Malone et al., 2009, Li et al., 2009), nor how piRNA biogenesis initiates in the absence of maternally inherited piRNAs, for example, during early mammalian embryogenesis (Aravin et al., 2008) or following horizontal transposon transfer.

Here we investigate piRNA specificity in the Drosophila ovary, which harbors two mechanistically distinct pathways: a slicer-competent, ping-pong-driven germline pathway and a slicer-independent somatic follicle cell pathway (Brennecke et al., 2007, Gunawardane et al., 2007, Malone et al., 2009, Li et al., 2009, Senti and Brennecke, 2010, Olivieri et al., 2010, Nishimasu et al., 2012, Voigt et al., 2012). By dissecting the molecular logic of both systems, we discover a unifying principle: piRNA biogenesis is founded on pervasive, largely indiscriminate sampling of cytoplasmic transcripts, with specificity emerging through tissue-specific modules that exploit intrinsic vulnerabilities of transposable elements.

## RESULTS

### Selective piRNA output within a slicer-independent piRNA pathway

Somatic follicle cells in the *Drosophila* ovary deploy a slicer-independent piRNA pathway in which nuclear Piwi silences the expression of endogenous retroviruses (Malone et al., 2009, Sienski et al., 2012, Rozhkov et al., 2013, Le Thomas et al., 2013). To study piRNA specificity without confounding germline input, we used cultured ovarian somatic stem cells (OSCs), which harbor a fully functional somatic piRNA pathway centered on nuclear Piwi (Fig. 1A,B) (Niki et al., 2006, Lau et al., 2009, Saito et al., 2009). High-complexity small RNA profiling of Argonaute-bound species (Fig. S1A) (Grentzinger et al., 2020, Jayaprakash et al., 2011) and mapping to a de novo OSC genome assembly (Handler and Brennecke, 2025) revealed that ∼65% of OSC piRNAs derive from transposon sequences with strong antisense bias (∼95%), nearly all mapping to the three active somatic piRNA clusters (*flamenco, 77B, 20A*). However, thousands of host mRNA 3′ UTRs also yield piRNAs, comprising ∼20% of the total pool (Fig. 1C,D) (Saito et al., 2009, Robine et al., 2009). These were quantified using a curated annotation pipeline that accounts for isoform-specific 3′ UTR usage and CDS invariance (Fig. S1B,C). All piRNAs shared hallmarks of Zucchini-dependent processing: 5′-uridine preference and robust phasing (Fig. S1D–F).

**Figure 1.**
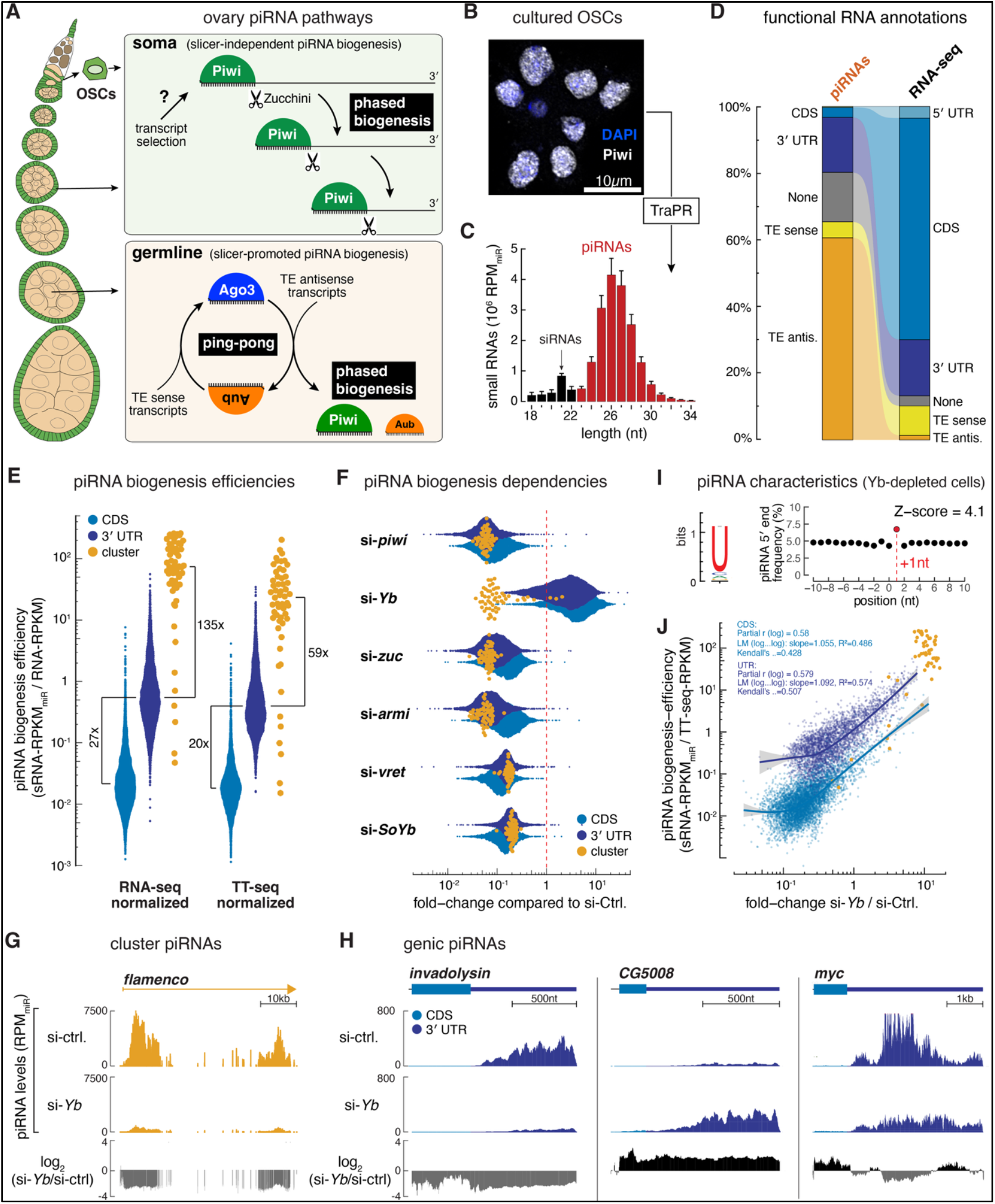
Yb globally determines piRNA biogenesis efficiency in the somatic pathway. **(A)** Schematic of a *Drosophila* ovariole illustrating piRNA biogenesis pathways in somatic cells (phased biogenesis only; exclusive pathway in OSCs) and germline cells (ping-pong amplification coupled with phased biogenesis). **(B)** Cultured, immortalized ovarian somatic cells (OSCs) expressing nuclear Piwi as the sole PIWI-clade protein (anti-Piwi immunofluorescence, white; DAPI, blue). **(C)** Length distribution (nucleotide) of TraPR-isolated (Argonaute-loaded) small RNAs in OSCs, normalized to miRNA abundance (reads per million miRNAs, RPMmiR). miRNAs are omitted; siRNAs are indicated; piRNAs (23– 35 nt) are shown in red; error bars reflect variation across biological replicates (n=5). **(D)** Fractional annotation of OSC piRNAs compared with total RNA-seq reads, partitioned into mRNA coding sequences (CDS), 5′ and 3′ untranslated regions (UTRs), transposable elements (sense and antisense), and unannotated regions (None). **(E)** piRNA biogenesis efficiencies (log_10_) for CDS (light blue), 3′ UTRs (dark blue), and piRNA cluster precursors (yellow), calculated as piRNA abundance normalized to precursor RNA levels (RNA-seq) or transcriptional output (TT-seq). Efficiencies for CDS and 3′ UTRs were computed per full-length unit, whereas cluster efficiencies were calculated using genome-unique 500-nt tiles. Data are shown as jittered dot plots with medians indicated by horizontal lines; fold differences are noted. **(F)** Fold changes (log_10_) in piRNA levels from CDS, 3′ UTRs, and piRNA clusters in OSCs depleted of the indicated piRNA biogenesis factors (Piwi, Yb, Zuc, Armi, Vret, SoYb) relative to control cells (si-Ctrl). Each dot represents a full-length CDS or 3′ UTR, or a 500-nt piRNA cluster tile. **(G)** Genome browser tracks showing piRNA profiles (RPM_miR_) across the first 50 kb of the *flamenco* piRNA cluster in control and Yb-depleted OSCs. The lower track displays the log_2_ fold change (si-*Yb*/si-ctrl). **(H)** Genome browser tracks for representative genic piRNA sources (*invadolysin, CG5008, myc*) in control and Yb-depleted OSCs, shown as piRNA abundance (RPM_miR_) and log2 fold-change (si-*Yb*/si-ctrl). **(I)** piRNA features in Yb-depleted OSCs. Left, sequence logo showing 5′ nucleotide bias (1U). Right, phasing analysis depicting the frequency of piRNA 5′ ends at positions −10 to +10 relative to piRNA 3′ ends on the same strand; the Z-score for the +1 position is indicated. **(J)** Relationship between piRNA biogenesis efficiency (E) and Yb dependency (F), both shown as log_10_ values, for piRNAs mapping to CDSs, 3′ UTRs, or 500-nt piRNA cluster tiles. Regression fits for CDS and 3′ UTR classes and associated statistics are shown.

Strikingly, different transcript classes varied dramatically in piRNA biogenesis efficiency (Fig. 1D; Fig. S1G). When piRNA levels per transcript were normalized to steady-state precursor abundance or transcriptional output, cluster RNAs generated on average ∼100-fold more piRNAs per kilobase than typical 3′ UTRs and >1,000-fold more than mRNA coding sequences (Fig. 1E). However, efficiencies overlapped substantially: some 3′ UTRs approached cluster-level output, while some cluster regions performed no better than average UTRs. This >1,000-fold range in processing efficiency across single-stranded precursors indicated that precursor selection must be a major determinant of piRNA output.

### Yb specifies piRNA precursor selection in the somatic pathway

To identify the genetic basis of this selectivity, we depleted established pathway factors by RNAi and quantified piRNA levels. Depletion of Piwi, Zucchini, Armitage, Vreteno, or SoYb caused uniform pan-piRNA loss, indicating that a shared machinery processes all precursor classes into Piwi-bound piRNAs (Fig. 1F) (Saito et al., 2010, Nishimasu et al., 2012, Zamparini et al., 2011, Ipsaro et al., 2012, Handler et al., 2013, Olivieri et al., 2010, Handler et al., 2011, Hirakata et al., 2019).

In striking contrast, depletion of the DEAD-box protein Yb (Olivieri et al., 2010, Saito et al., 2010) produced a highly non-uniform phenotype (Fig. 1F) (Ishizu et al., 2019, Ishizu et al., 2015): Cluster-derived piRNAs, including those from *flamenco*, were reduced to levels comparable to those observed upon Zucchini or Armitage knockdown, consistent with derepression of Piwi-controlled LTR retrotransposons (Fig. 1F,G and Fig. S1H). Genic piRNAs instead exhibited transcript-specific responses—some were strongly reduced, others unaffected, and most increased, with changes often varying even within individual transcripts (Fig. 1F,H) (Ishizu et al., 2019, Hirakata et al., 2019). Importantly, remaining piRNAs retained Zucchini-dependent processing signatures (Fig. 1I; Fig. S1I-K), indicating that Yb functions upstream of core biogenesis machinery by influencing precursor selection (Ishizu et al., 2019).

Notably, Yb dependency correlated strongly with intrinsic processing efficiency: highly productive precursors required Yb, whereas low-output precursors increased upon Yb depletion (Fig. 1J). This analysis also confirmed that coding sequences consistently yield fewer piRNAs than 3′ UTRs, a difference we investigate further below.

To test whether these findings extend in vivo, we analyzed Piwi-bound piRNAs from somatic follicle cells of ovaries. The somatic piRNA profile resembled that of OSCs, with antisense transposon piRNAs predominating and genic piRNAs, mainly from 3′ UTRs, making up ∼10% (Fig. S1L). Yb depletion by transgenic RNAi strongly reduced piRNAs from the *flamenco* cluster, while 3′ UTR piRNAs from soma-specific transcripts showed transcript-specific responses similar to those observed in OSCs (Fig. S1M). Accordingly, transcripts exhibiting Yb-dependent piRNA production in OSCs also required Yb for efficient processing in vivo (Fig. S1N). Altogether, these findings established Yb as a global specifier of piRNA precursor competence in the somatic pathway.

### Uridine-rich sequences license Yb-dependent piRNA biogenesis

To identify RNA features promoting piRNA processing, we focused on mRNA 3′ UTRs, which generate abundant piRNAs yet lack the mapping ambiguities inherent to transposon sequences. Because piRNA output varied within individual transcripts (Fig. 1H), we subdivided mRNA 3′ UTRs into 100-nt tiles and quantified piRNA biogenesis efficiency for each segment, normalizing for transcript abundance (see methods). Efficiencies spanned >1,000-fold (Fig. 2A).

**Figure 2.**
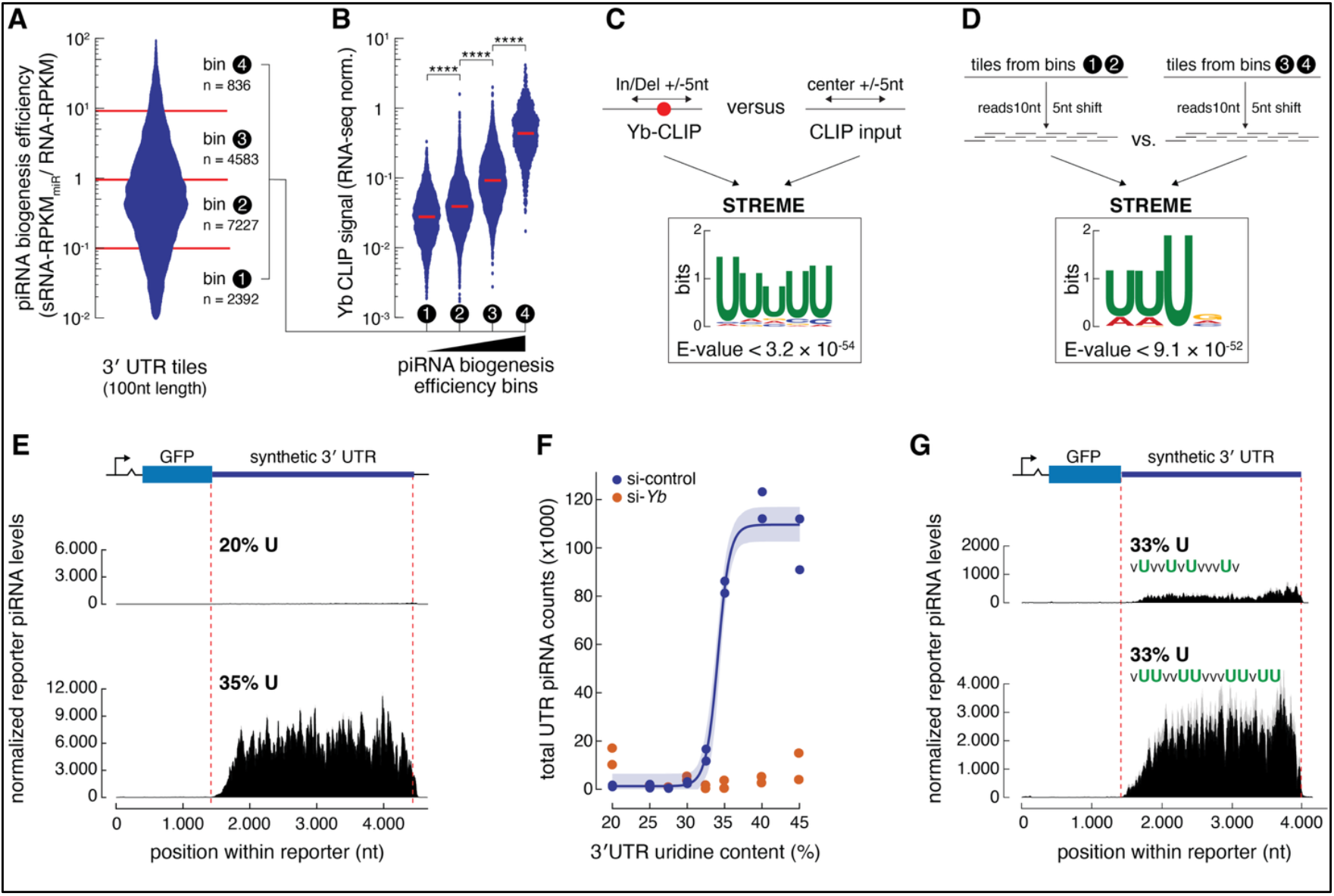
The uridine content of an RNA defines Yb’s impact on piRNA output. **(A)** Distribution of piRNA biogenesis efficiencies of 100-nt 3′ UTR tiles, calculated as piRNA abundance normalized to precursor RNA levels. Tiles were grouped into four efficiency bins (n as indicated). **(B)** Yb CLIP-seq signal (normalized to RNA-seq) for 3′ UTR tiles grouped by piRNA biogenesis efficiency as in (A); red bars denote medians. ****, *p* < 0.0001. **(C)** De novo motif discovery at Yb CLIP sites using STREME. Sequences centered on crosslink-associated indels (±5 nt) from Yb CLIP libraries were compared with matched windows from CLIP input libraries (±5 nt). **(D)** De novo motif discovery using STREME comparing low-efficiency (bins 1–2) vs. high-efficiency (bins 3–4) 3′ UTR tiles, using 10-nt windows with a 5-nt offset as shown. **(E)** GFP reporter constructs with synthetic 3′ UTRs of defined uridine content. piRNA profiles from low-U (20% U) and high-U (35% U) reporters are shown, normalized to 1M miRNAs and corrected for reporter RNA abundance. Mean of two biological replicates; shading shows min–max range. **(F)** Reporter piRNA output, normalized to 1M miRNAs and corrected for reporter RNA abundance, plotted against synthetic 3′ UTR uridine content in control (black dots) and Yb-depleted (red dots). Each dot represents one biological replicate (two replicates per sensor construct and U-content). **(G)** GFP reporters with equal uridine content (33% U) but different U configurations (V = A, G, or C). Shown are piRNA profiles for dispersed U versus clustered UU motifs, normalized to reporter RNA abundance. Lines represent the mean across biological replicates; shading indicates the min–max range.

To test whether this broad dynamic range correlates with Yb binding, we performed Yb CLIP-seq. Yb occupancy correlated strongly with biogenesis efficiency, with highly productive tiles showing robust Yb enrichment (Fig. 2B; Fig. S2A) (Ishizu et al., 2019).

De novo motif discovery from CLIP reads containing crosslink-induced signatures identified a uridine-rich motif as the most significantly enriched Yb-binding element (Fig. 2C). Strikingly, an independent comparison of productive versus non-productive tiles—performed without using CLIP data—likewise identified U-rich sequences as the top predictor of piRNA output (Fig. 2D). Even total uridine content alone correlated strongly with piRNA production, whereas other nucleotides showed no comparable association (Fig. S2B).

Previous studies had proposed that Yb recognizes a specific, structured cis-regulatory element within the *traffic jam* (*tj*) 3′ UTR (Ishizu et al., 2015, Ishizu et al., 2019, Takase et al., 2022). To evaluate whether such sequence-specific recognition underlies Yb-dependent piRNA production, we generated clonal OSC lines expressing GFP reporters carrying different versions of the *tj* 3′ UTR from a defined genomic landing site. To distinguish reporter from endogenous *tj* piRNAs, we introduced discriminatory SNPs. The SNP-modified *tj* UTR generated piRNA levels comparable to the endogenous gene (Fig. S2C). However, shuffling the previously reported Yb-binding site—which disrupts its proposed secondary structure—had no detectable effect on piRNA output (Fig. S2C). These results suggested that *tj*’s pronounced piRNA production is not determined by a specific structural motif but may instead reflect broader sequence features. Consistent with this interpretation, the *tj* 3′ UTR exhibits elevated uridine content (32%).

To directly test whether U-content alone is sufficient to confer precursor competence, we generated OSC lines expressing GFP reporters with entirely synthetic 3′ UTRs spanning 20–45% uridine, with remaining nucleotides equally distributed among A, G, and C (Fig. 2E,F; Fig. S2D). piRNA production scaled dramatically with U-content: low-U reporters (20–28%) produced minimal piRNAs, while high-U reporters (35–45%) generated approximately 100-fold more, with robust production emerging around 32% U (Fig. 2E,F; Fig. S2E). Critically, this U-dependent response was Yb-dependent: upon Yb depletion, high-U reporters lost the majority of their piRNAs while low-U reporters gained piRNAs, flattening the U-content/output relationship (Fig. 2F; Fig. S2F–H).

To explore which U-configurations are most effectively recognized, we constructed reporters with identical UTR length (∼2.5 kb) and U-content (∼33.5%) but with uridines arranged as isolated U, UU, or UUU motifs. Despite identical nucleotide composition, reporters containing UU or UUU motifs produced approximately 10-fold more piRNAs than those with only single uridines (Fig. 2G; Fig. S2I), identifying the UU dinucleotide as a critical Yb-recognized feature.

Together, these experiments establish that Yb selectively routes transcripts into piRNA biogenesis by recognizing U-rich sequences, particularly UU dinucleotides. Although Zucchini cleavage and Piwi loading both favor uridines (Izumi et al., 2020, Stein et al., 2019), our data demonstrate that Yb, not the core machinery, imposes U-dependent precursor selection: upon Yb depletion, U-poor sequences gain piRNAs while U-rich sequences lose them, opposing responses that are incompatible with intrinsic machinery bias but consistent with Yb steering U-rich precursors into piRNA processing.

### U-rich initiation sites propagate piRNA production downstream

Phased piRNA biogenesis proceeds from 5′ to 3′ (Mohn et al., 2015, Han et al., 2015, Homolka et al., 2015). To investigate how local U-content influences output across larger regions, we embedded defined U-rich or U-poor segments within constant-sequence neighborhoods of transgenes. We first inserted a 2kb sequence from *Drosophila yakuba* into the center of a synthetic 3′ UTR in both orientations, creating reporters in which the internal region had either 40% or 24% U-content, while the surrounding sequence maintained a uniform 33% U-content (Fig. 3A). In the high-U configuration, piRNA production peaked within the central fragment, consistent with its elevated U-content relative to the flanking regions. In the low-U configuration, piRNA output collapsed within the central region; however, the decrease did not occur immediately at the boundary. Instead, piRNA levels declined gradually over ∼1.5 kb in the 5′→3′ direction. Notably, the central fragment strongly affected output from the immediately adjacent sequences, despite those sequences being identical. This demonstrates that sequence content imposes a “neighbourhood effect” that influences processing efficiency, especially downstream.

**Figure 3.**
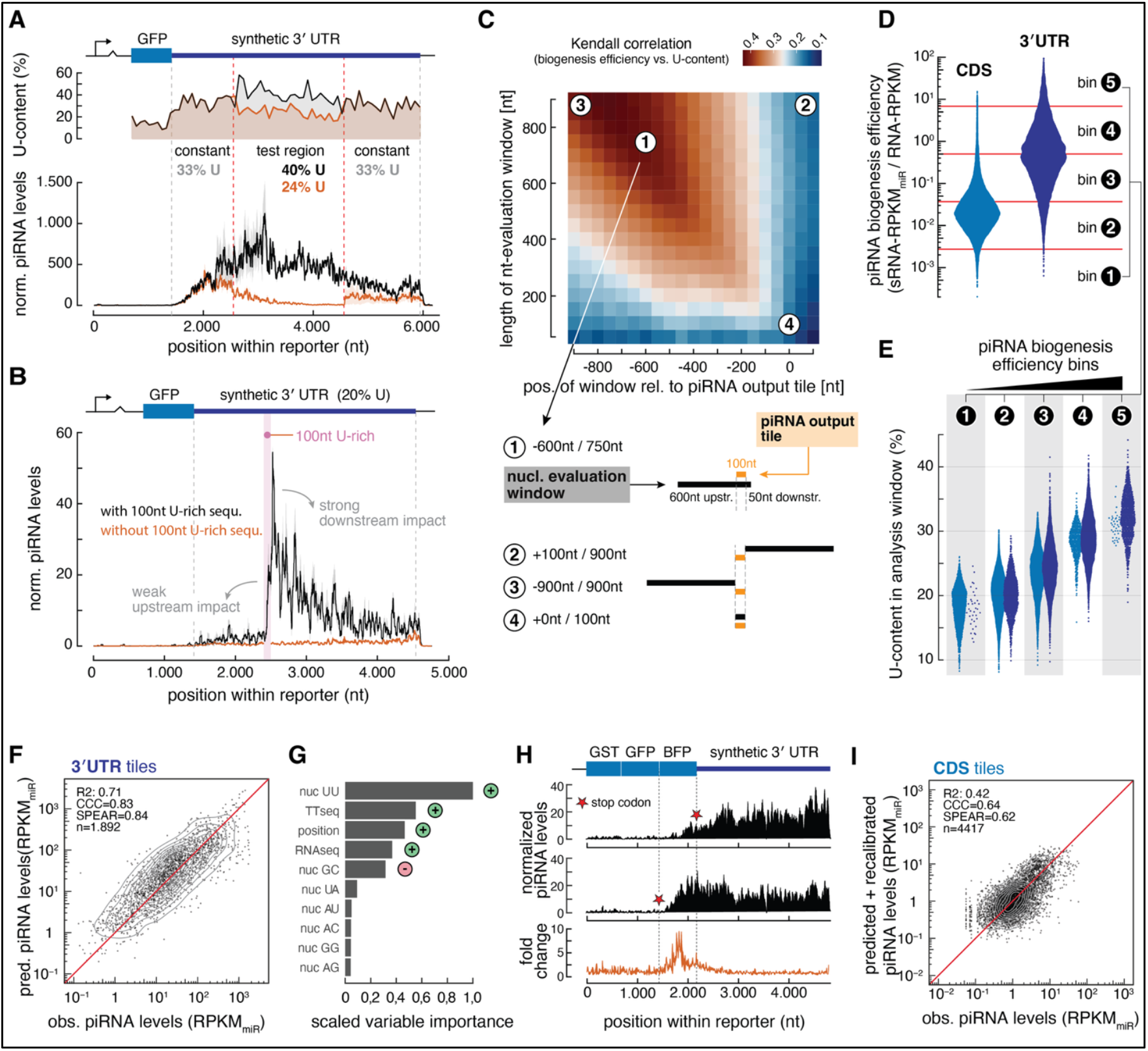
Quantitative piRNA output prediction from U-content and biogenesis directionality. **(A)** GFP reporter constructs with 3′ UTRs featuring indicated U-content profiles with constant flanking (∼33%) and variable central (∼24% versus ∼40%) regions. Shown below are corresponding piRNA profiles for low central U (black) and high central U (∼35%, orange) reporters, normalized to 1M miRNAs and corrected for reporter RNA abundance. Data represent the mean of two biological replicates; shading indicates the min–max range. **(B)** As in (A), but for reporters with U-poor synthetic 3′ UTRs (20% U) containing (black) or lacking (orange) a central 100-nt U-rich segment (33%). **(C)** Heat map showing Kendall correlations between local piRNA biogenesis efficiency and U-content in evaluation windows of varying length and position relative to a 100-nt piRNA output tile; example window configurations (1 to 4) are marked and annotated below. **(D)** Distribution of piRNA biogenesis efficiencies (log_10_) for 100-nt CDS tiles and 100-nt 3′ UTR tiles, computed as piRNA levels normalized to precursor abundance (y-axis: sRNA-RPKM_miR_ / RNA-RPKM) and grouped into five efficiency bins. **(E)** U-content of the analysis window (750nt long, shifted -600nt) associated with 3′ UTR tiles grouped into the five piRNA biogenesis efficiency bins from (D). **(F)** Comparison of predicted and observed piRNA levels for test-set 3′ UTR tiles (n = 1892) from a gradient-boosting model trained on sequence composition, RNA abundance (RNA-seq), transcriptional output (TT-seq), and positional features. Performance metrics are indicated. **(G)** Scaled variable importance from the 3′ UTR prediction model. The +/-symbols indicate if a variable positively or negatively impacts piRNA levels. **(H)** Same as (A), using reporters assessing the impact of translation on piRNA output from a coding sequence. Normalized piRNA profiles for translated and non-translated BFP ORFs (stop codon indicated) followed by a synthetic 3′ UTR are shown; bottom panel shows the position-resolved fold-change (non-translated/translated). **(I)** Predicted (after recalibration) versus observed piRNA levels for CDS tiles using the 3′ UTR–trained model; performance metrics are indicated.

To resolve this effect at higher resolution, we inserted a 100-nt U-rich segment (33% U) into an otherwise U-poor (20% U), 3-kb 3′ UTR that alone produced negligible piRNAs (Fig. 3B). The U-rich element increased piRNA production by several dozen-fold. piRNA levels rose across the U-rich segment, peaked just downstream, and decayed over 1.5–2 kb, demonstrating that U-poor sequences can generate substantial piRNAs when positioned downstream of U-rich regions.

### Quantitative prediction of piRNA output from 3′ UTR sequence features

Based on our reporter assays, high local U-content boosts piRNA output of a downstream sequence. To test whether this principle extends to endogenous transcripts, we systematically scanned mRNA 3′ UTRs to identify the window size and position whose U-content best predicts piRNA production for each 100-nt tile (Fig. 3C). Windows larger than ∼600 nt yielded the strongest correlations, but only when positioned asymmetrically: most of the informative sequence lay upstream of the focal tile. UU-dinucleotide frequency was equally predictive, whereas no other mono- or dinucleotide feature exhibited comparable predictive power (Fig. S3A,B). Thus, endogenous piRNA output reflects U-density integrated over extended upstream regions (Fig. 3D,E; Fig. S3C).

To ask whether nucleotide content is sufficient to quantitatively predict piRNA output across 3′ UTRs. We trained a gradient boosting machine model using piRNA levels per 100-nt tile as the response variable. Predictors included di-nucleotide composition within the asymmetric window, steady-state RNA abundance (RNA-seq), nascent transcriptional output (TT-seq), tile position within the UTR, and UTR length. The model achieved high predictive accuracy (statistical values) on held-out data (Fig. 3F).

Feature importance analysis showed that UU-dinucleotide enrichment and RNA availability (TT-seq and RNA-seq) were the strongest predictors of piRNA output (Fig. 3G). Tile position also contributed, reflecting a ∼500-nt “ramp-up” region of piRNA output at the start of 3′ UTRs (Fig. S3D) (Robine et al., 2009). This pattern is consistent with phased processing, in which local and transcript-internal initiation events within the 3′ UTR generate a 5′→3′ series of piRNAs whose output gradually decays. When many such events occur across a transcript population, their trailing products overlap, producing the observed gradual increase.

Thus, piRNA output from mRNA 3′ UTRs is determined by U-dependent internal initiation combined with transcript abundance, providing a quantitative framework that explains the wide dynamic range of endogenous piRNA profiles.

### Translation antagonizes phased piRNA biogenesis

Consistent with their low U-content (average 21%), most mRNA coding sequences produced minimal piRNAs that increased upon Yb depletion (Fig. 1E,F; Fig. 3D) (Robine et al., 2009, Ishizu et al., 2019). The rare high-output CDS tiles instead were unusually U-rich (Fig. 3E) and Yb-dependent (Fig. 1J).

When applied to CDSs, the 3′ UTR-trained model accurately captured relative differences in piRNA output, spanning ∼100-fold across tiles (Fig. S3E). However, the model systematically overestimated absolute piRNA output from CDSs by nearly an order of magnitude, suggesting CDS-specific inhibition. Hypothesizing that translation suppresses piRNA production, we generated a reporter in which the Azurite BFP coding sequence was either translated or rendered non-translated by repositioning the stop codon. Although the U-rich Azurite CDS produced piRNAs even when translated, shifting the stop codon upstream by 700 nucleotides increased piRNA output from the same sequence by up to 8-fold (Fig. 3H).

Thus, translation antagonizes phased piRNA biogenesis independent of sequence content. After correcting for this offset, the model accurately predicted CDS-derived piRNAs (Fig. 3I).

### A common uridine-recognition logic unifies cluster and mRNA piRNA biogenesis

The *flamenco* piRNA cluster—the archetypal somatic cluster spanning ∼700 kb of densely packed antisense LTR retrotransposon fragments (Goriaux et al., 2014b, Zanni et al., 2013, Sarot et al., 2004, Prud’homme et al., 1995, Handler and Brennecke, 2025)—provides a critical test: if Yb-dependent U-recognition governs somatic piRNA production, even *flamenco* should obey these sequence rules rather than requiring cluster-specific licensing.

*flamenco* is transcribed from a single RNA polymerase II promoter (Goriaux et al., 2014a, Mohn et al., 2014), but transcription is not uniform across the locus, declining by several hundred-fold along its length (Handler and Brennecke, 2025). Using genome-unique regions within *flamenco* as anchors, we estimated local transcription rates and steady-state RNA levels to calculate piRNA biogenesis efficiency for cluster-unique 100-nt tiles (Fig. S4A, B).

*flamenco* tiles exhibited ∼45-fold higher efficiency than average 3′ UTRs and ∼800-fold higher than CDS tiles (Fig. 4A), an advantage largely lost upon Yb depletion, with output falling to near-UTR levels (Fig. 4B). Consistent with U-driven processing, local U-content within *flamenco* strongly correlated with piRNA production, with the most prolific tiles exceeding 40% U or 15% UU (Fig. 4C; Fig. S4C).

**Figure 4.**
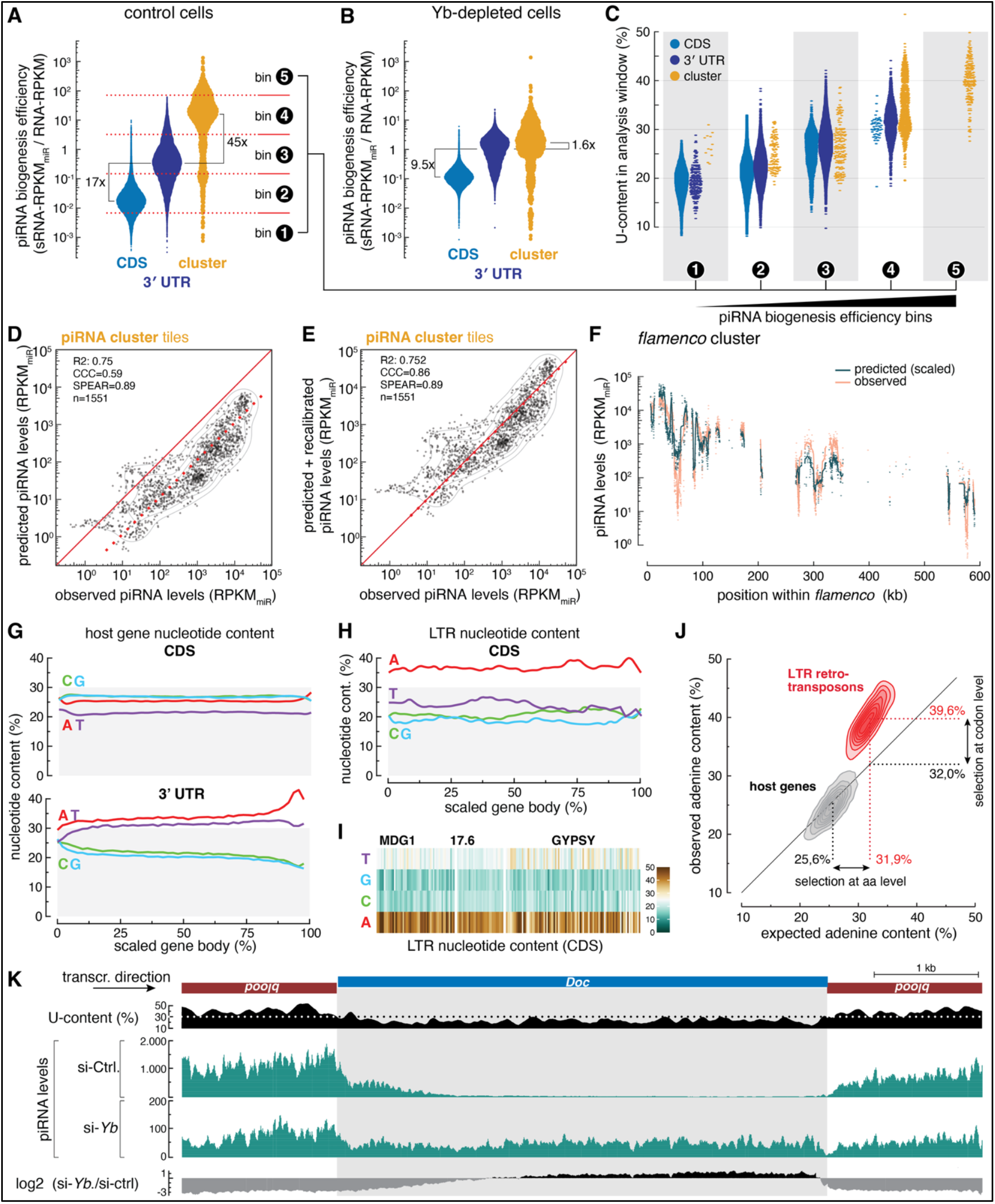
Adenosine bias in retrotransposon genomes accounts for *flamenco*’s exceptional piRNA output. **(A)** piRNA biogenesis efficiency (piRNA-RPKM_miR_ / RNA-RPKM) of 100-nt CDS, 3′ UTR, and piRNA cluster tiles in control OSCs, stratified into five efficiency bins. **(B)** As in (A) but in Yb-depleted OSCs. **(C)** U-content of the analysis window associated with 100nt CDS, 3′ UTR, and piRNA cluster tiles grouped by piRNA biogenesis efficiency (bins 1–5). **(D)** Comparison of predicted and observed piRNA levels for cluster tiles using the 3′ UTR–trained model (Fig. 3). Performance metrics are indicated. **(E)** As in (D) but after recalibration. **(F)** Observed and predicted (recalibrated) piRNA levels (log_10_) along the *flamenco* cluster plotted as a function of genomic position. **(G)** Average nucleotide content along *Drosophila melanogaster* genic coding sequences (CDS) and 3′ UTRs plotted along scaled gene body position. **(H)** Average nucleotide content along LTR retrotransposon CDS plotted along scaled transposon body position. **(I)** Heat map showing nucleotide composition across coding sequences of selected LTR retrotransposon families among Drosophilid species (Repbase entries from the MDG1, 17.6, and GYPSY groups). **(J)** Expected (based on codon usage in *D. melanogaster*) versus observed adenosine content for *D. melanogaster* host genes (grey) and Drosophilid LTR retrotransposons (red; Repbase entries from MDG1, 17.6, and GYPSY groups), with arrows indicating contributions from skewed amino-acid composition (horizontal shift) and synonymous codon usage (vertical shift). **(K)** UCSC genome browser tracks depicting a region in *flamenco* with a nested *Doc* sense insertion within a *blood* antisense insertion. Shown are local U-content, piRNA levels in control and Yb-depleted OSCs, and log_2_ fold changes in piRNA levels (si-*Yb*/si-control) across the region.

To test whether the same sequence-based rules govern piRNA production from *flamenco* and mRNAs, we applied the 3′ UTR–trained predictive model to the cluster. The model explained the ∼1,000-fold variation in piRNA output across the locus (R^2^ ≈ 0.75) but underpredicted absolute output (Fig. 4D). This offset might reflect the competing activity of translation—present on mRNAs but absent from *flamenco*. After correction, predicted and observed piRNA levels aligned closely across the entire locus (Concordance correlation coefficient [CCC] = 0.82) (Fig. 4E,F). Removing nucleotide composition as a model feature markedly reduced predictive power (R^2^ = 0.31; CCC = 0.50) (Fig. S4D).

Thus, the same Yb-dependent U-recognition mechanism governs both cluster and mRNA piRNA biogenesis. This convergence raised a fundamental question: how does a simple, broadly acting U-recognition mechanism nevertheless achieve transposon specificity?

### Adenosine-rich retrotransposon sequences generate U-rich *flamenco* transcripts

The *flamenco* cluster is densely populated by antisense LTR retrotransposon fragments (Brennecke et al., 2007, Zanni et al., 2013, Handler and Brennecke, 2025). Based on this organization, we hypothesized that *flamenco*’s U-richness directly reflects the intrinsic nucleotide composition of its resident transposons. To test this idea, we compared the nucleotide content of host mRNAs with that of *flamenco*-resident LTR families from all structurally intact Repbase entries in Drosophilid species (*17*.*6, gypsy, mdg1*) (Zanni et al., 2013, Senti et al., 2025) (Fig.4G-I).

This analysis revealed a remarkable contrast. Whereas host mRNAs are GC-rich in their coding regions and AU-rich in their 3′ UTRs, LTR retrotransposon sequences, coding as well as non-coding, show exceptionally high adenosine (A) levels, frequently exceeding 40%. When these A-rich elements are transcribed in the antisense direction—as they are in *flamenco*—their A-bias is converted into U-rich RNA. This observation raised two questions: what molecular features of retrotransposon sequences account for their extreme A-content, and does transposon nucleotide bias indeed underlie *flamenco*’s pronounced responsiveness to Yb?

We first analyzed how LTR retrotransposon ORFs achieve their elevated A-content (Lerat et al., 2002). Using average *Drosophila melanogaster* codon usage as a reference (Vicario et al., 2007), we compared expected versus observed A-content across host coding regions and retrotransposon ORFs (*gag* and *pol*) (Fig. 4J). Two features distinguished LTR sequences. First, for any given amino acid, transposons preferentially use A-rich synonymous codons, raising their observed A-content to ∼39% compared with the ∼32% predicted from host codon usage (Fig. S4E). Second, retrotransposon proteins are biased toward amino acids whose codons are A-rich. While the amino-acid composition of an average host protein requires codons with an expected A-content of ∼26%, transposon proteins require ∼32% (Fig. 4J). For example, isoleucine and lysine (encoded by A-rich codons) are enriched in transposon proteins, whereas the respectively similar residues leucine and arginine (encoded by A-poor codons), are depleted (Fig. S4F).

If transposon A-richness underlies *flamenco*’s Yb-dependent piRNA production, local orientation flips of transposon insertions should create position-specific differences in piRNA output. We focused on a naturally occurring “orientation switch” in *flamenco* where a 5kb sense *Doc* insertion (U-content ∼22.5%) is flanked by two antisense *blood* fragments (U-content ∼40%) (Fig. 4K). The *blood* fragments rank among the highest-efficiency piRNA sources genome-wide. At the antisense→sense boundary, piRNA production declined gradually over ∼1.5 kb into the U-poor *Doc* region, precisely as seen for our reporter constructs. At the sense→antisense boundary, piRNA output rose steeply within a few hundred nucleotides, reaching the high-efficiency regime characteristic of U-rich regions. Thus, U-rich sequences within *flamenco* act as potent piRNA sources, whereas adjacent U-poor sequences do not. Critically, Yb depletion abolished the sense– antisense asymmetry: *blood* piRNAs dropped >10-fold while *Doc* piRNAs slightly increased (Fig. 4K).

These findings fundamentally challenge the cluster-centric model. *flamenco*’s exceptional piRNA productivity does not arise from cluster-specific licensing but from an extreme manifestation of nucleotide composition bias encoded in retrotransposon genomes themselves. By preferentially engaging U-rich regions, Yb naturally selects antisense transposon fragments, converting a simple biochemical preference into sequence-specific transposon targeting.

### Basal transcript sampling forms the basis of the somatic piRNA pathway

Unlike core factors whose depletion causes near-complete piRNA collapse, Yb-depleted OSCs retained 48% ± 8% of total piRNA levels (Fig. 5A), with dramatically altered composition: mRNAs accounted for ∼70% of piRNAs versus ∼20% in wild-type cells (Fig. 5B), and cluster processing efficiency relative to median UTRs dropped from ∼50-fold to <2-fold (Fig. 4A,B). This redistribution suggested that in Yb’s absence, a basal biogenesis machinery samples cytoplasmic RNAs largely indiscriminately, with Yb functioning as a specificity filter that steers this basal program toward U-rich transcripts (see also (Ishizu et al., 2019, Ge et al., 2019)).

**Figure 5.**
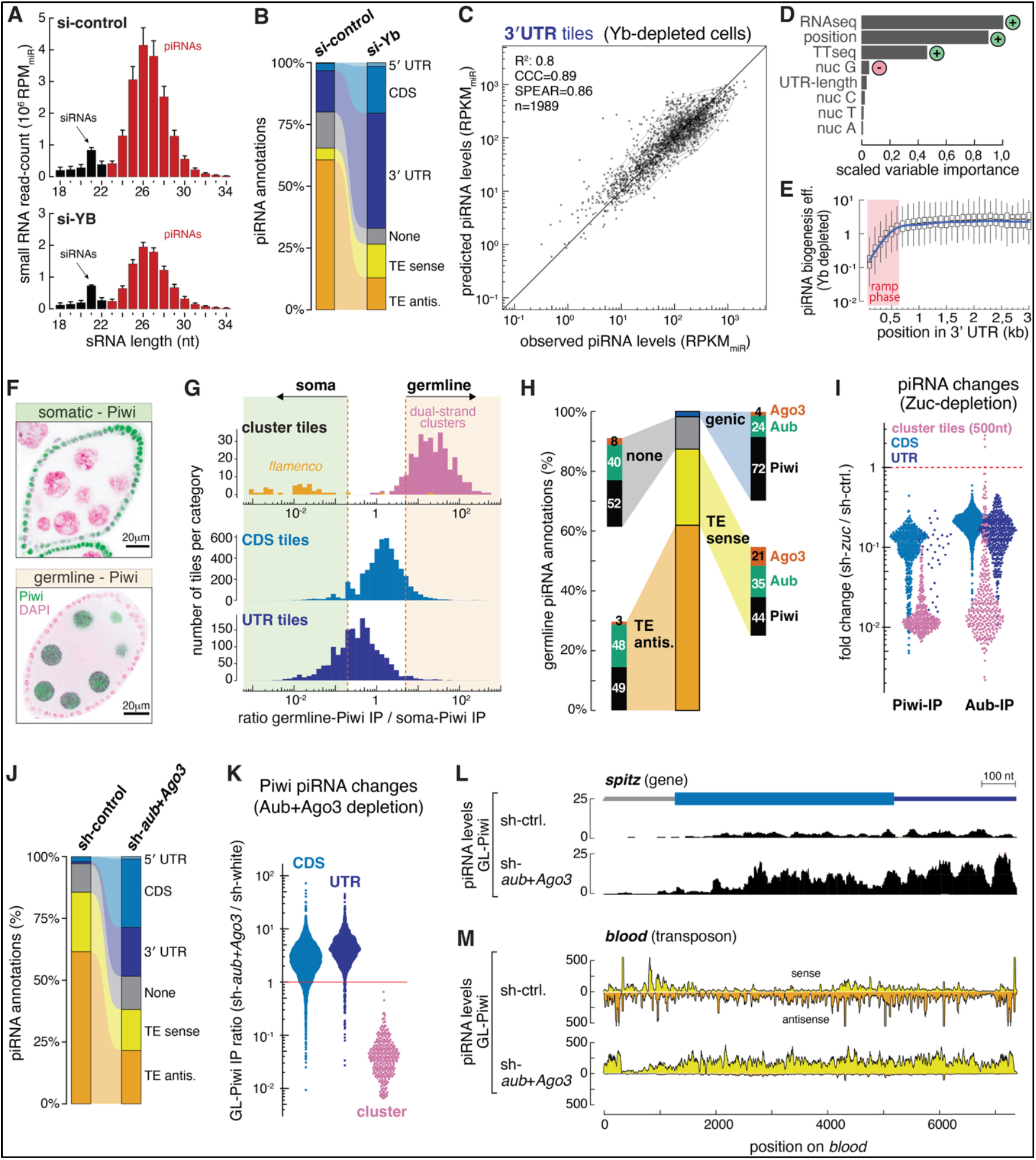
Naïve transcript sampling forms the basis of piRNA biogenesis in soma and germline. **(A)** Length distribution (nucleotides) of TraPR-isolated (Argonaute-loaded) small RNAs from control and Yb-depleted OSCs, normalized to miRNA abundance (reads per million miRNAs, RPM_miR_). miRNAs are omitted; siRNAs are indicated; piRNAs (23–35 nt) are shown in red. Error bars represent variation across biological replicates (control, *n* = 5; si-Yb, *n* = 4). **(B)** Genomic origin of OSC piRNAs partitioned into coding sequences (CDS), 3′ untranslated regions (3′ UTRs), transposable elements (TEs; sense and antisense), and unannotated regions (None) under control and Yb-depleted conditions. **(C)** Predicted versus observed piRNA abundance from 3′ UTR tiles in Yb-depleted cells, using a gradient-boosting model trained on Yb-depleted data. Model performance metrics are indicated. **(D)** Scaled variable importance from the Yb-depleted 3′ UTR prediction model. The +/-symbols indicate if a variable positively or negatively impacts piRNA levels. **(E)** Distribution of piRNA biogenesis efficiencies (piRNA output normalized for transcriptional output - TT-seq) per 100nt tile across 3′ UTRs in Yb-depleted cells, with the early “ramp” region in red. **(F)** Representative confocal images of egg chambers expressing GFP-Piwi specifically in somatic (top) or germline cells (bottom) used for soma Piwi-IP and germline Piwi-IP libraries. **(G)** Distributions of the germline Piwi-IP to soma Piwi-IP ratio (log10) for piRNAs derived from CDS, 3′ UTR, and cluster annotations (*flamenco* and dual-strand germline clusters); dashed lines indicate ≥5-fold enrichment in the germline or soma Piwi-IP libraries. **(H)** Annotation-based composition of germline piRNA sources from a composite library of Piwi-, Aub-, and Ago3-bound small RNAs (center). Flanking stacked bar plots indicate the distribution of each piRNA population among the three PIWI proteins. **(I)** Log10 fold changes in Piwi- and Aub-bound piRNAs following Zucchini depletion in the germline, shown separately for CDS-, 3′ UTR–, and dual-strand cluster–derived piRNAs. Full-length CDS and 3′ UTR features were quantified, whereas dual-strand clusters were partitioned into 500-nt blocks for quantification. **(J)** Annotation-based composition of germline Piwi-bound piRNAs in control and Aub + Ago3–depleted ovaries. **(K)** Log_10_ fold changes in germline Piwi-bound piRNAs upon Aub+Ago3 depletion, shown as Piwi-IP ratios (sh-*aub*+*Ago3*/sh-*white*) for CDS-, 3′ UTR-, and dual-strand cluster–derived piRNAs. **(L)** Representative gene locus (*spi*) showing germline Piwi-bound piRNA levels (RPM_miR_) in sh-control and sh-*aub*+*Ago3* ovaries. **(M)** Profile of germline Piwi-bound piRNAs mapping to the *blood* LTR retrotransposon in sh-control and sh-*aub*+*Ago3* ovaries (sense piRNAs in yellow, antisense in orange).

To characterize basal sampling, we trained a prediction model on piRNA data from 100-nt 3′ UTR tiles in Yb-depleted OSCs. Predictor variables included nucleotide content, transcriptional output (TT-seq), steady-state RNA levels (RNA-seq), UTR length, and tile position within the UTR. The model achieved high predictive accuracy on 100-nt 3′ UTR tiles from an independent set of test genes (Fig. 5C).

Variable importance analysis revealed that in the absence of Yb, nucleotide composition exerts no influence on biogenesis efficiency (Fig. 5D). Instead, RNA availability (RNA-seq and TT-seq) together with tile position within the UTR emerged as the dominant determinants. As in control cells, piRNA output increased gradually in the first ∼500 nucleotides of 3′ UTRs (Fig. 5E), and coding sequences produced ∼ tenfold fewer piRNAs than 3′ UTRs, indicating this inhibition operates independently of Yb (Fig. 4A,B). To assess whether RNA availability alone determines piRNA output, we excluded the first 600 nucleotides of 3′ UTRs from the prediction. A model incorporating only TT-seq and RNA-seq features provided a good prediction of piRNA abundance (R^2^ = 0.62; CCC = 0.806; Fig. S5A,B).

Thus, the somatic piRNA pathway comprises a naïve biogenesis system that samples transcripts according to availability, with Yb layering U-dependent specificity onto this inherently non-selective foundation.

### Germline piRNA specificity emerges from basal sampling coupled to ping-pong amplification

The somatic pathway achieves specificity through Yb-dependent U-recognition. But germline cells, the principal site of transposon and piRNA pathway activity, do not express Yb (Qi et al., 2010, Olivieri et al., 2010, Saito et al., 2010). Instead, germline piRNA populations are shaped by cytoplasmic PIWI proteins with slicer activity. Cleavage of target transcripts by Aubergine (Aub) or Ago3 fuels ping-pong amplification—a self-reinforcing cycle that enriches complementary piRNA pairs and triggers phased biogenesis loading Piwi and Aub (Brennecke et al., 2007, Gunawardane et al., 2007, Mohn et al., 2015, Han et al., 2015, Homolka et al., 2015). However, while ping-pong defines mature germline populations, it remains unclear how piRNA identity is initially established absent pre-existing piRNAs.

The prevailing model assigns this initiating role to piRNA cluster transcripts, proposing that antisense transposon sequences embedded within clusters give rise to primary piRNAs that subsequently engage active transposon RNAs in ping-pong. However, our discovery of basal transcript sampling in somatic cells suggested an alternative framework: naïve piRNAs may arise through largely indiscriminate cytoplasmic sampling, with specificity emerging only during ping-pong amplification when complementary transcripts are encountered, a property effectively restricted to transposon sequences. In this model, clusters supply antisense relay transcripts rather than conferring initial specificity.

To test whether basal transcript sampling operates in germline cells, we first examined Piwi-bound piRNAs, which are generated by phased, Zucchini-dependent processing in both soma and germline. Comparing Piwi-associated piRNAs isolated from somatic follicle cells versus germline revealed expected partitioning into soma-specific loci (e.g., *flamenco*) and germline-specific loci (dual-strand piRNA clusters) (Mohn et al., 2014, Senti et al., 2015) (Fig. 5F,G). In addition, more than a thousand host mRNAs (CDSs and 3′ UTRs) produced Piwi-bound piRNAs, and these showed soma-specific, germline-specific, and mixed origin (Fig. 5G). Basal generation of genic piRNAs is therefore a feature of both pathways.

Comprehensive profiling of piRNAs bound to Piwi, Aub, or Ago3 in germline cells revealed that mRNA-derived piRNAs comprise approximately 2% of the total germline pool and load preferentially into Piwi and Aub (Fig. 5H). These genic piRNAs derive almost exclusively from sense strands, display robust phasing, and lack ping-pong signatures—features paralleling the basal population in Yb-depleted soma (Fig. S5C,D). Notably, germline cells generate proportionally more CDS-derived piRNAs than somatic cells, potentially reflecting higher levels of translationally repressed mRNAs, thereby making CDSs available for processing (Fig.S5E).

To test whether germline genic piRNAs arise through phased processing, we depleted Zucchini specifically in germline cells. This caused near-complete loss of cluster-derived piRNAs and several-fold reductions in Piwi- and Aub-bound genic piRNAs (Fig. 5I). Residual genic piRNAs retained a phased signature (Fig. S5F), suggesting incomplete Zucchini depletion or minor contribution from alternative biogenesis pathways. Thus, Zucchini-dependent processing generates basal piRNAs in germline as in soma.

If basal sampling provides an indiscriminate piRNA foundation, removing ping-pong amplification should elevate this underlying population. We analyzed Piwi-bound piRNAs in the germline upon depletion of the slicer-competent proteins Aub and Ago3. Under these conditions, ping-pong was eliminated and total Piwi piRNA levels dropped approximately fourfold (Senti et al., 2015). The compositional shift of Piwi-bound piRNAs was dramatic: whereas transposon-derived piRNAs dominate wild-type germline Piwi complexes (with genic piRNAs at ∼3%), genic piRNAs comprised nearly 50% of Piwi-bound species in ping-pong-deficient ovaries (Fig. 5J). Consistent with this shift, cluster-derived piRNAs collapsed more than 20-fold, while genic piRNAs—retaining phased signatures (Fig. S5G)—increased two-to fourfold (Fig. 5K; Fig. 5L shows a representative example).

Loss of ping-pong also revealed basal sampling for transposon sense transcripts. For example, the LTR element *blood*, which in wildtype ovaries displays mixed sense and antisense Piwi-bound piRNAs, consistent with slicer-induced phased biogenesis, switched to an almost complete sense piRNA profile covering the entire element (Fig. 5M). Similar shifts to sense piRNA populations were evident for several other transposons (Fig. S5H).

These results establish that germline piRNA specificity arises from ping-pong amplification operating atop a basal biogenesis system that samples transcripts with limited intrinsic selectivity. Initial, low-level piRNA generation from active transposon transcripts establishes a basal pool. Upon encountering complementary antisense transcripts, an inevitable consequence of transposon mobilization into host genes, these naïve piRNAs trigger ping-pong amplification. Thus, the pathway exploits a universal vulnerability of mobile elements: their mobility itself generates the antisense transcripts required for adaptive, sequence-specific silencing.

## DISCUSSION

We have identified a unifying principle underlying transposon-specific piRNA production: the pathway is founded on pervasive, largely indiscriminate sampling of cytoplasmic transcripts, with sequence specificity emerging through tissue-specific mechanisms that exploit intrinsic, population-level properties of transposable elements. Rather than relying on hard-coded genomic loci or element-specific recognition, the system converts fundamental features of mobile elements, their atypical nucleotide composition and their mobility, into selective piRNA production.

In the germline, naïve transcript sampling coupled to catalytically active Argonaute proteins exploits a core vulnerability of transposable elements: their inevitable production of antisense RNAs following mobilization into host transcription units. Low-level piRNA generation from active transposon transcripts establishes a basal pool of sense piRNAs. Upon expression of complementary antisense transcripts, these naïve piRNAs engage ping-pong amplification, an inherently sequence-specific mechanism that selectively enriches piRNAs against active elements. The presence of phased genic piRNAs in germline Piwi and Aub complexes, and their increased abundance in ping-pong-deficient ovaries, demonstrates that adaptive amplification builds upon indiscriminate basal biogenesis.

Because naïve piRNAs are generated at low levels, they compete with established ping-pong pairs for biogenesis machinery access, providing a mechanistic explanation for the observation that acquisition of piRNA-mediated immunity against newly invading transposons requires multiple generations (Jensen et al., 1999, Josse et al., 2007, Khurana et al., 2011, Kofler et al., 2015). In this context, maternal inheritance of PIWI–piRNA complexes (Blumenstiel and Hartl, 2005, Brennecke et al., 2008, Le Thomas et al., 2014) ensures that newly established sequence-specific responses are transmitted to subsequent generations, enabling gradual reinforcement of silencing.

The principle of naïve transcript sampling appears to act broadly. In fetal mouse testes, piRNA biogenesis initiates through broad sampling of host mRNAs and transposon sense RNAs in a Zucchini/Armitage-dependent, ping-pong-independent process (Aravin et al., 2008, Zheng et al., 2010, De Fazio et al., 2011, Watanabe et al., 2011, Pandey et al., 2013). Slicer-mediated amplification engages upon emergence of complementary transcripts, directing production of transposon antisense piRNAs (Aravin et al., 2008, De Fazio et al., 2011). Natural *KoRV-A* endogenization in koalas parallels this: basal, sense-biased piRNA production transitions to robust ping-pong amplification following antisense insertion into an expressed host gene (Yu et al., 2019, Yu et al., 2025).

In Drosophila somatic follicle cells, which lack slicer-competent PIWI proteins and ping-pong amplification, specificity is achieved through a distinct but equally parsimonious mechanism. The DEAD-box protein Yb selectively promotes piRNA biogenesis from uridine-rich RNA precursors. Because antisense RNAs derived from LTR retrotransposons are intrinsically U-rich—a direct consequence of the unusually high adenosine content of retrotransposon genomes—this mechanism preferentially channels transposon-complementary transcripts into the piRNA pathway. Recognition of such a minimal nucleotide feature implies a multivalent mode of Yb action. Homotypic interactions mediated by the Yb N-terminal Hel-C domain (Hirakata et al., 2019) may amplify weak sequence biases across extended transcript regions, providing a mechanism for integrating U-content over larger sequence regions. Through this compositional mechanism, Yb can enhance piRNA production by up to two orders of magnitude, independent of transcript identity.

The *flamenco* piRNA cluster exemplifies this principle at scale. Composed almost entirely of antisense LTR retrotransposon fragments, *flamenco* transcripts are non-coding and exceptionally U-rich, rendering them highly efficient piRNA precursors. These findings necessitate reconceptualizing piRNA clusters. Rather than uniquely licensed loci, clusters function primarily as sources of transposon antisense transcripts—optimal Yb substrates in soma due to U-richness, ping-pong relay elements in the germline. In both contexts, orientation is decisive: antisense insertions efficiently drive piRNA production, whereas sense insertions fail to do so. Recent demonstrations that antisense insertions outside canonical clusters elicit robust piRNA-mediated silencing reinforce this view (Konstantinidou et al., 2024, Yu et al., 2025, Rafanel et al., 2025).

The exploitation of transposon adenosine-richness for somatic piRNA specificity aligns with an emerging paradigm: hosts discriminate self from non-self by reading population-level sequence properties. Mammalian HUSH targets A-rich, intron-poor nascent RNAs, silencing LINEs and endogenous retroviruses through heterochromatin (Seczynska et al., 2022, Lehner, 2025). SAF-B family RNA-binding proteins exploit the A-richness of transposon sequences to suppress aberrant splice site usage arising from intronic LINE insertions (Ilik et al., 2024). Thus, transposon A-content constitutes a broadly conserved non-self signal independently recognized by distinct surveillance pathways.

Zooming out, our results establish naïve transcriptome sampling as a foundational principle of the piRNA pathway across tissues. In both germline and soma, basal piRNA biogenesis depends on the conserved factors Zucchini and Armitage, with Armitage likely serving as a point of integration for tissue-specific specificity mechanisms, potentially through regulation of its ATPase activity (Ge et al., 2019, Ishizu et al., 2019, Rogers et al., 2017). In an accompanying study, Tomari and colleagues independently discover naïve piRNA biogenesis and demonstrate that it is conserved in insects and mammals, reinforcing the generality of our findings (Shoji and Tomari, 2026). This modular architecture, coupling non-specific sampling to context-dependent filters, enables flexible, adaptive genome defense against continually emerging threats. By converting intrinsic vulnerabilities of mobile elements into selective triggers, the piRNA pathway exemplifies how biological surveillance systems can achieve robust self/non-self discrimination without exhaustive genetic encoding—a principle shared by immune systems, quality control pathways, and other systems confronting diverse, evolving challenges.

## ACKNOWLEDGEMENTS

We thank the VBCF and IMBA/IMP/GMI core facilities for support, particularly the NGS facility for sequencing, the BioOptics unit for support, and the VDRC for fly stocks. The computational results presented were obtained using the CLIP cluster (https://clip.science). Members of the Brennecke laboratory and alumni (P. Andersen, L. Baumgartner, J. Schnabl-Baumgartner) gave valuable comments on the manuscript. This work was supported by the Austrian Academy of Sciences, a European Research Council advanced grant (ERC-AdG-101142075), and the Austrian Science Fund (FWF grant P 36970). A.T is supported by a DOC Fellowship from the Austrian Academy of Science. For the purpose of open access, the authors have applied a CC BY public copyright license to any Author Accepted Manuscript version arising from this submission.

## MATERIALS AND DATA AVAILABILITY

Fly stocks used in this study are available from the VDRC (https://shop.vbc.ac.at/vdrc_store). The sequencing data generated for this study have been deposited in the NCBI Gene Expression Omnibus (GEO) under the accession number ##### (pending GEO processing due to US government shutdown). Published datasets used in this manuscript include small RNA and RNA sequencing of studies GSE213383, GSE64802, GSE71775. A detailed inventory of all datasets, including published data, is provided in Supplementary table S2.

The computational code is deposited at GitHub https://github.com/BrenneckeLab/Handler_2026-YB.

## DECLARATION OF GENERATIVE AI AND AI-ASSISTED TECHNOLOGIES

During the preparation of this work, the authors used Abacus AI to polish writing, as well as to assist in the development of analysis scripts. After using this tool, the authors reviewed and edited the content as needed and take full responsibility for the content of the publication.

## AUTHOR CONTRIBUTIONS

Conceptualization: DH, RH, JB

Methodology: DH, RH

Investigation: DH, RH, WYWW, AT

Data Curation: DH

Validation: DH

Formal analysis: DH, RH

Visualization: DH, RH, JB

Resources: JB

Funding acquisition: JB

Project administration: JB

Supervision: JB

Writing – original draft: DH, JB

Writing – review & editing: DH, RH, JB

## List of Supplementary Materials

Figures S1-S5

Materials and Methods

Tables S1-S3

