## Supplementary Figures and Methods for "A Naïve RNA Sampling Core Enables Adaptive piRNA Specificity Against Transposable Elements"

**The Supplemental materials include:**

Figs. S1 to S5
Materials and Methods
Table S1
Supplementary References

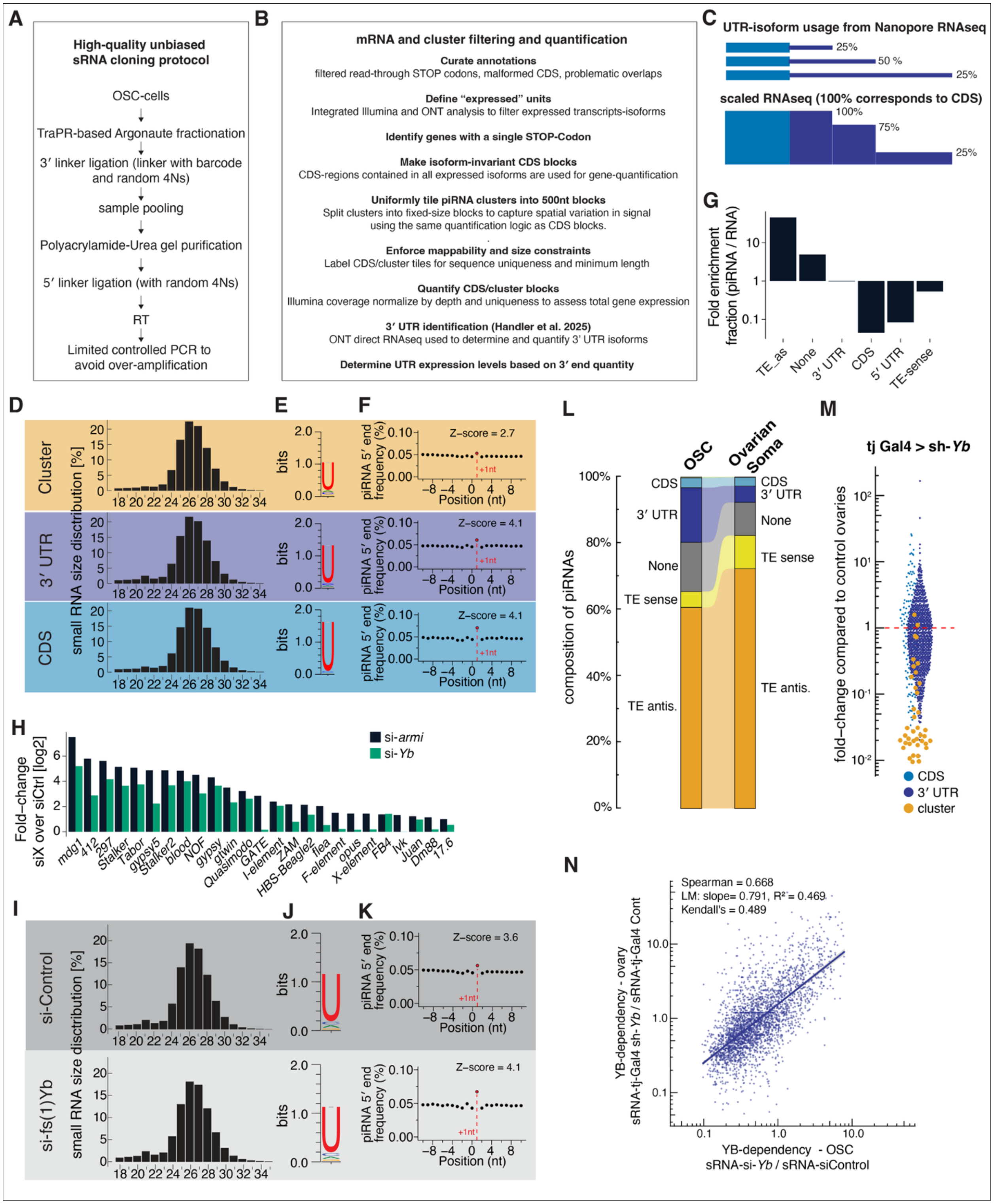


**Figure S1. Related to Figure 1**

**(A)** Schematic of the small RNA cloning workflow from ovarian somatic cells (OSCs), including TraPR-based purification of Argonaute-bound small RNAs and low-bias library preparation.

**(B)** Overview of the filtering/curation pipeline used for transcript annotation and quantification of genic features and piRNA cluster output, including definition of isoform-invariant CDS units and fixed-size cluster tiles.
**(C)** Schematic illustrating inference of 3′ UTR isoform usage from Nanopore (ONT) RNA-seq and the corresponding scaling of Illumina RNA-seq gene quantification using CDS coverage extended across the entire 3′ UTR (see Methods).
**(D–F)** piRNA features (separate for piRNAs mapping to CDS, 3′ UTR, cluster) in control OSCs. (E), small RNA length profile. (F), sequence logo showing 5′ nucleotide bias (1U). (G), phasing analysis depicting the frequency of piRNA 5′ ends at positions −10 to +10 relative to piRNA 3′ ends on the same strand; the Z-score for the +1 position is indicated.
**(G)** Fold enrichment ($log_{10}$) of annotation categories in piRNA populations relative to RNA-seq, calculated as the ratio of the fraction of small RNA-seq reads to the corresponding fraction of RNA-seq reads for each category (annotation classes as in Fig. 1D).
**(H)** Fold-change (log2) of transposon-derived RNA relative to control upon depletion of Armitage (si-*armi*) or Yb (si-*Yb*), shown for transposons increased ≥2× upon Armitage depletion.
**(I–K)** As in (D–F), for all piRNAs in control and Yb-depleted OSCs.
**(L)** Composition of piRNAs by annotation class in OSCs an in ovarian soma.
**(M)** Fold changes ($log_{10}$) in Piwi-bound piRNA levels from CDS, 3′ UTRs, and piRNA clusters (analyzed are soma-specific mRNAs and cluster tiles) in ovaries depleted of Yb in the soma (tj-Gal4 > sh-Yb) relative to control ovaries (si-ctrl). Each dot represents a full-length CDS or 3′ UTR, or a 500-nt piRNA cluster tile.
**(N)** Comparison of Yb dependency of mRNA-derived piRNAs (soma-specific 3′ UTRs) between OSCs and ovaries (Piwi-IP), plotted as $log_{10}$ fold change in piRNA levels upon Yb depletion in OSCs versus ovarian soma. Correlation statistics are indicated.

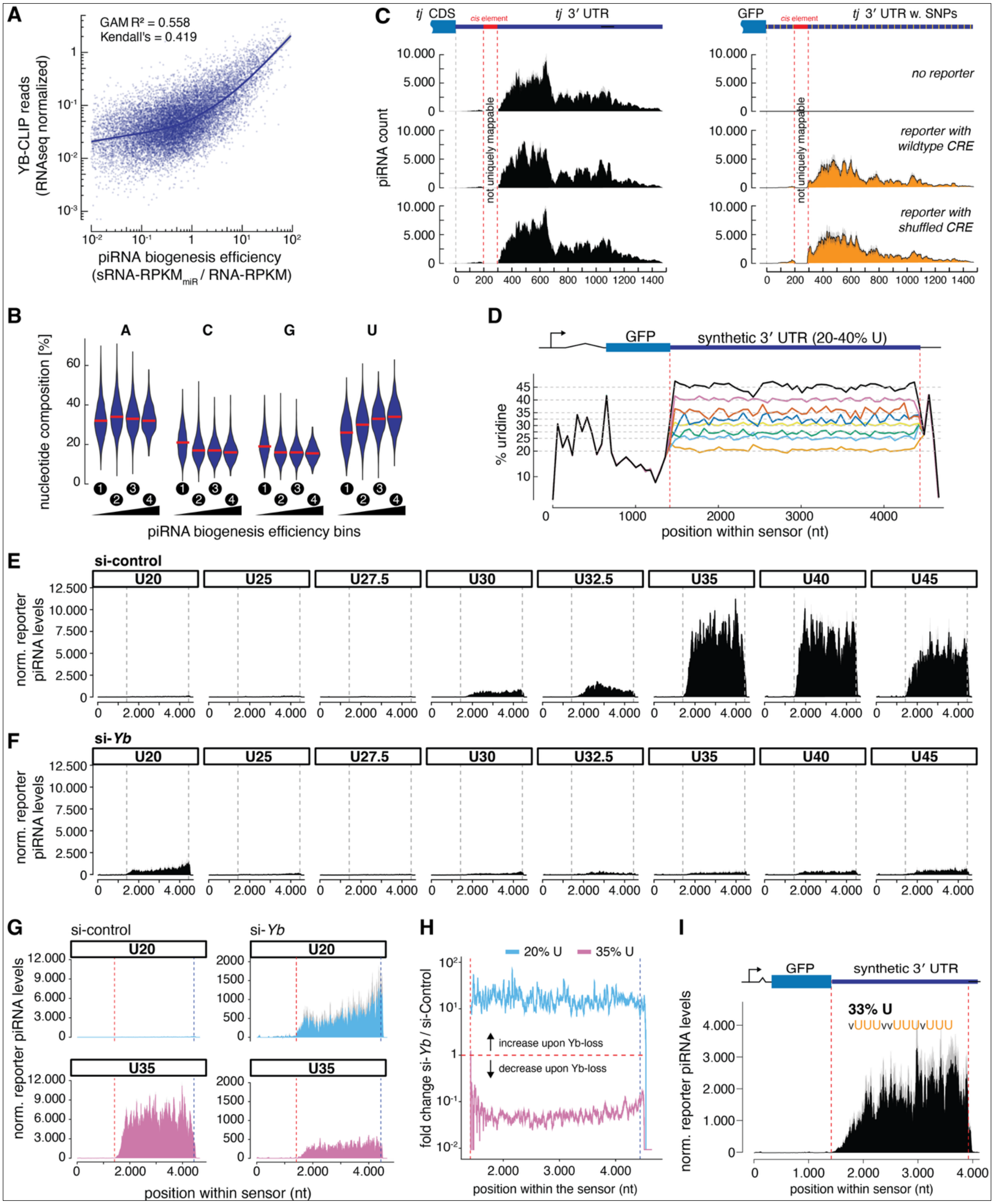


**Figure S2. Related to Figure 2**

**(A)** Yb CLIP-seq signal (RNA-seq–normalized) versus piRNA biogenesis efficiency (piRNA-$RPKM_{miR}$ / RNA-RPKM) for 100-nt 3′ UTR tiles. Fit and correlation statistics are indicated; both axes are $log_{10}$.

**(B)** Violin plots of A, C, G, and U content in 3′ UTR tiles stratified by piRNA biogenesis efficiency bins; red bars denote medians.
**(C)** piRNA profiles mapping to the endogenous *tj* mRNA (left) or the piRNA reporter with a *tj* 3′ UTR that carries discriminatory SNPs (right) in OSCs harboring no reporter (top), the reporter with the wildtype *tj* cis-element (middle) or the shuffled *tj* cis-element (bottom).
**(D)** Synthetic 3′ UTR reporter design with uridine content ranging 20–45% and remaining nucleotides equally distributed among A, C, and G. Position-resolved %U along the transcription unit is shown.
**(E)** piRNA profiles across the transcription unit for the U-content series (U20–U45%) in si-ctrl conditions, with 3′ UTR boundaries indicated. piRNA levels are normalized to reporter RNA levels. Data represent the mean of two biological replicates.
**(F)** As in (D) but in OSCs depleted of Yb (si-*Yb*).
**(G)** Representative piRNA profiles for U20 and U35 reporters in si-ctrl and si-Yb conditions (different scale). Data show the mean of two biological replicates.
**(H)** Position-resolved $\log_{10}$ fold-change in reporter piRNA levels upon Yb depletion (si-*Yb*/si-ctrl) for representative low-U (20% U) and high-U (35% U) reporters.
**(I)** piRNA profile of a reporter containing clustered UUU motifs (overall U-content 33%), with 3′ UTR boundaries marked. piRNA levels were normalized to reporter RNA levels. Data represent the mean of two biological replicates.

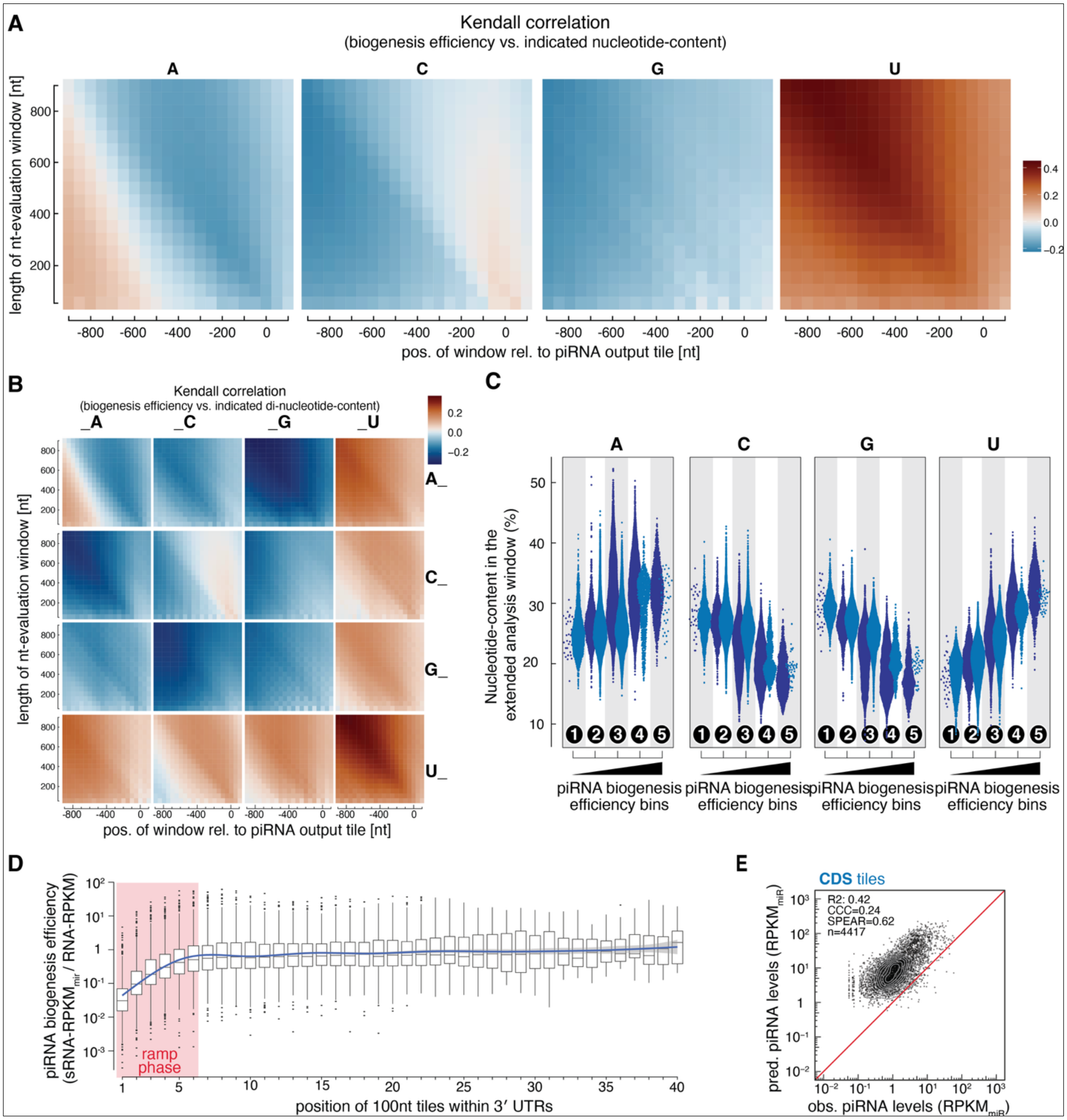


**Figure S3. Related to Figure 3.**

**(A)** Heat map showing Kendall correlations between local piRNA biogenesis efficiency and nt-content (A, C, G, U) in evaluation windows of varying length and position relative to a 100-nt piRNA output tile. Data for U identical as in to Figure 3C but with different color scheme.

**(B)** Heat map showing Kendall correlations between local piRNA biogenesis efficiency and UU dinucleotide content in evaluation windows of varying length and position relative to a 100-nt piRNA output tile. Directly comparable to panel (A) and Figure 3C.

**(C)** Average nt-content (A, C, G, U) of the analysis window (750nt long, shifted -600nt) associated with 3′ UTR tiles grouped into the five piRNA biogenesis efficiency bins from Figure 3D.

**(D)** piRNA biogenesis efficiencies across 3′ UTR sequences as a function of 100-nt tile position. Boxplots summarize tiles at each index. piRNA biogenesis efficiencies across 3′ UTR sequences as a function of 100-nt tile position. Boxplots summarize the distribution of efficiencies for tiles at each index. A smoothed trend line with shaded confidence interval indicates the overall trajectory of piRNA biogenesis efficiency along the 3′ UTR. The early "ramp" region of gradually increasing efficiency is marked.
**(E)** Predicted versus observed piRNA levels for CDS tiles using the 3′ UTR–trained model (raw output prior to recalibration); performance metrics are indicated.

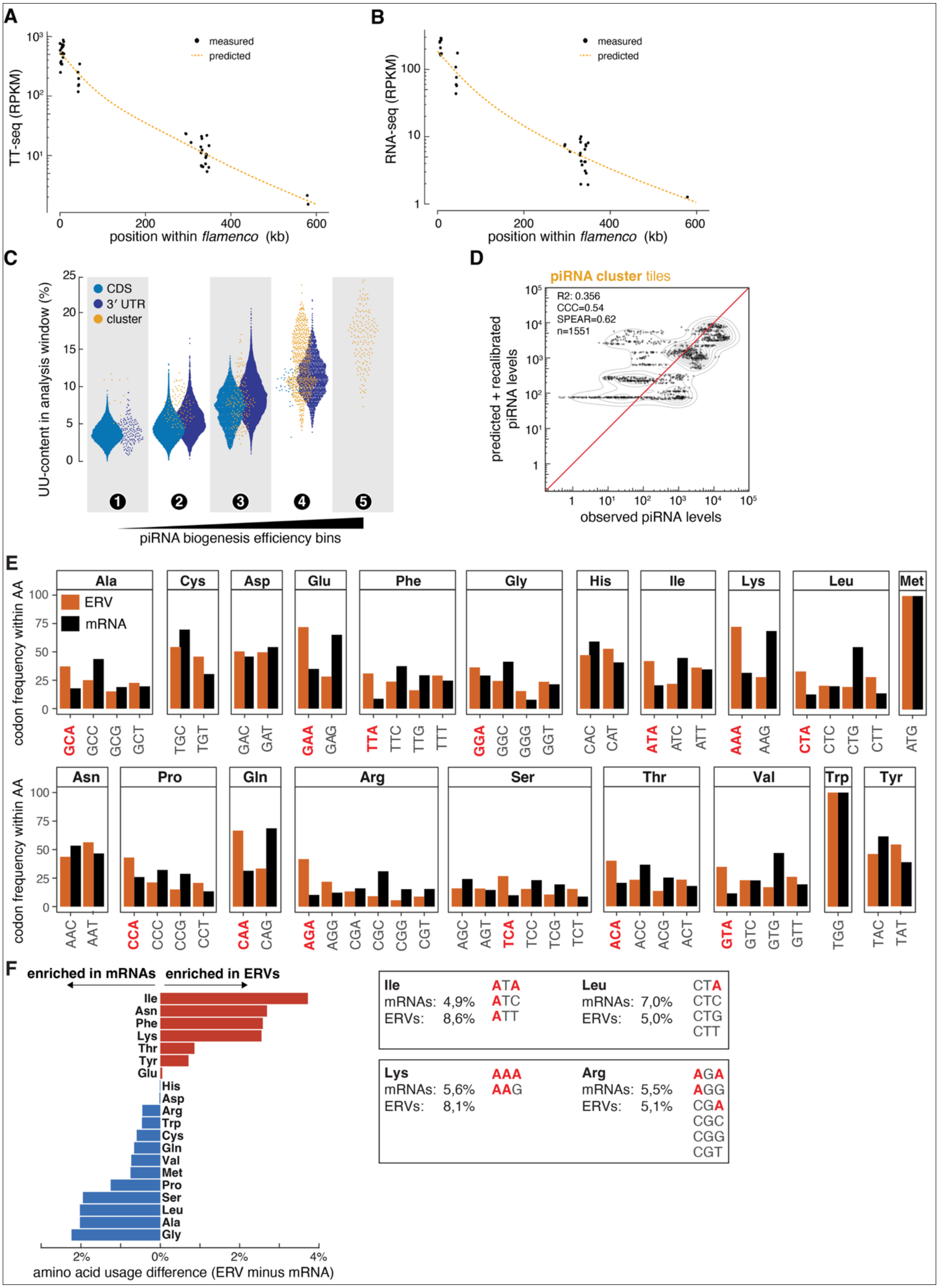
A
measured
predicted
TT-seq (RPKM)
position within flamenco (kb)
B
measured
predicted
RNA-seq (RPKM)
position within flamenco (kb)
C
CDS
3′ UTR
cluster
UU-content in analysis window (%)
piRNA biogenesis efficiency bins
D
piRNA cluster tiles
R2: 0.356
CCC=0.54
SPEAR=0.62
n=1551
predicted + recalibrated piRNA levels
observed piRNA levels
E
ERV
mRNA
codon frequency within AA
Ala Cys Asp Glu Phe Gly His Ile Lys Leu Met
Asn Pro Gln Arg Ser Thr Val Trp Tyr
F
enriched in mRNAs
enriched in ERVs
amino acid usage difference (ERV minus mRNA)
Ile mRNAs: 4,9% ERVs: 8,6% ATA ATC ATT
Leu mRNAs: 7,0% ERVs: 5,0% CTA CTC CTG CTT
Lys mRNAs: 5,6% ERVs: 8,1% AAA AAG
Arg mRNAs: 5,5% ERVs: 5,1% AGA AGG CGA CGC CGG CGT

**Figure S4. Related to Figure 4.**

**(A)** TT-seq signal (transcriptional output) across the *flamenco* cluster plotted by genomic position. Dots represent TT-seq RPKM values for 500-nt tiles with sufficient unique mappability; the predicted trend is shown as line.

**(B)** As in (A) but with RNA-seq signal.

**(C)** UU dinucleotide content within the analysis window associated with 100-nt CDS, 3′ UTR, and piRNA cluster tiles grouped into piRNA biogenesis efficiency bins (1–5).

**(D)** Predicted (after recalibration) versus observed piRNA levels for piRNA cluster tiles using the 3′ UTR–trained model but excluding all nucleotide composition features; performance metrics are indicated.

**(E)** Comparison of codon usage in *Drosophila melanogaster* endogenous genes (black bars) and insect endogenous retroviruses (orange bars). Per amino acid, the alternative codons with additional adenosine residues are indicated in red.

**(F)** Differences in amino acid usage between endogenous *D. melanogaster* genes and insect endogenous retroviruses (iERVs). Red bars indicate increased usage in iERVs, blue bars reduced usage. To the right, the amino acid usage of two pairs of chemically similar amino acids (Ile and Leu; Lys and Arg) in host genes and iERVs, respectively. Also shown are the possible codons for the amino acids, with adenosines colored in red.

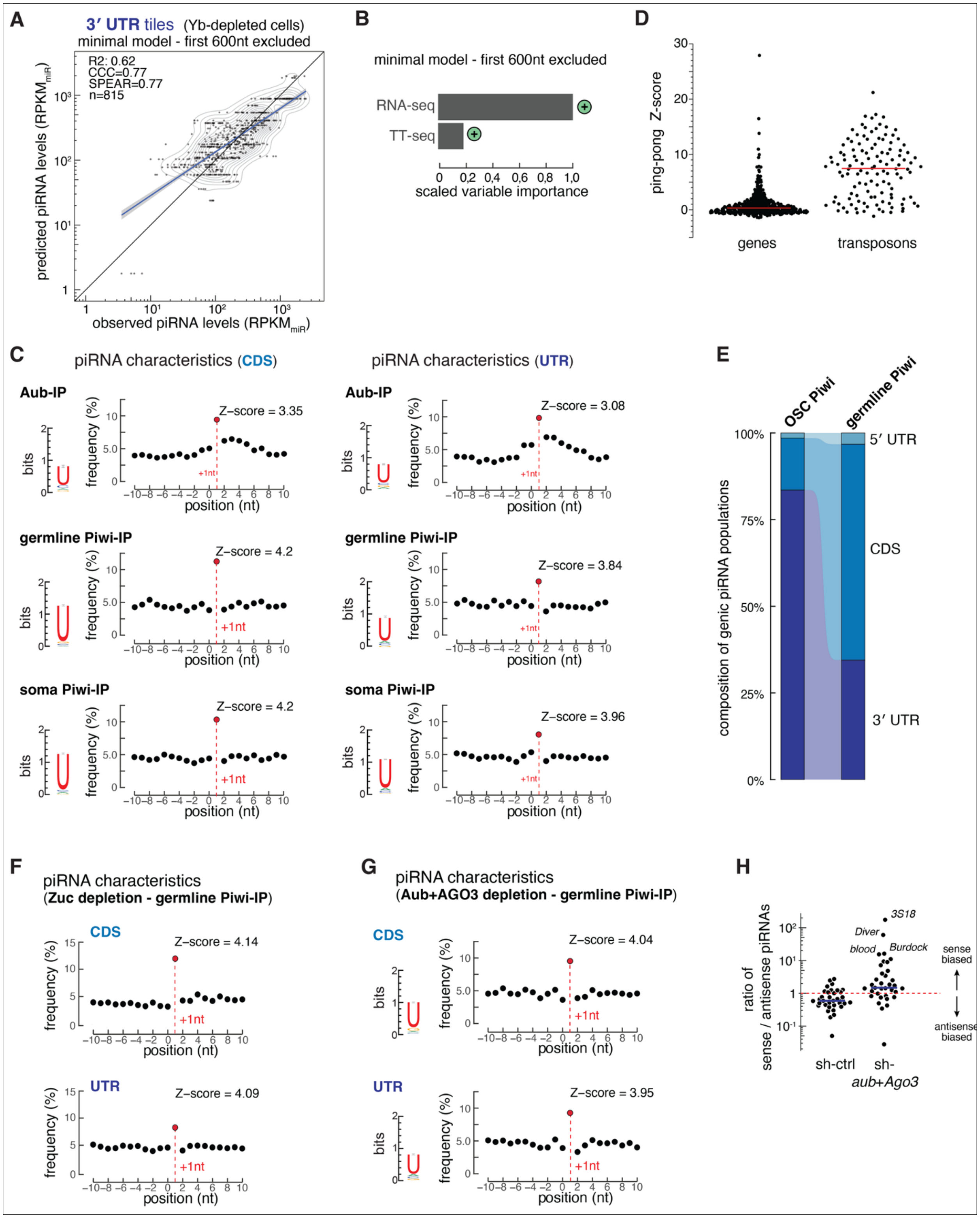


**Figure S5. Related to Figure 5.**

**(A)** Predicted versus observed piRNA abundance from 3′ UTR tiles in Yb-depleted cells using a minimal model excluding the first 600 nt of each 3′ UTR. Model performance metrics are indicated.

**(B)** Scaled feature importance from the minimal model in (A), highlighting contributions of RNA-seq and TT-seq–derived features. Both features are positively associated with piRNA abundance.
**(C)** Characteristics of CDS- and 3′ UTR–derived piRNAs in Aub immunoprecipitation (Aub-IP), germline Piwi-IP, and somatic Piwi-IP libraries. Shown are 5′ nucleotide bias (1U) and phasing profiles (−10 to +10 nt), depicting the frequency of piRNA 5′ ends relative to piRNA 3′ ends; Z-scores for +1 phasing enrichment are indicated.
**(D)** Distribution of ping-pong Z-scores calculated from germline piRNA populations, comparing genic piRNAs (little or no evidence of ping-pong amplification for most genes) with transposon-derived piRNAs (strong ping-pong signatures for most elements).
**(E)** Annotation-based composition of genic piRNAs in Piwi isolated from OSCs versus ovarian germline cells.
**(F)** piRNA characteristics for CDS- and 3′ UTR-derived piRNAs in germline Piwi-IP upon Zucchini depletion. Shown are 5′ nucleotide bias (1U) and phasing profiles (−10 to +10 nt), depicting the frequency of piRNA 5′ ends relative to piRNA 3′ ends; Z-scores for +1 phasing enrichment are indicated.
**(G)** as in (F) but for germline Piwi-IP piRNAs upon Aub+Ago3 depletion.
**(H)** Sense-to-antisense ratio of germline Piwi-bound piRNAs in sh-control and sh-*aub* + *Ago3* ovaries.

### MATERIALS AND METHODS

#### OSC cell culture

Ovarian somatic cells (OSCs) were maintained at 27 °C in a humidified incubator with 2.5% $CO_2$ in M3 medium supplemented with 10% fetal bovine serum (FBS), insulin, glutathione, and fly extract as previously described (Niki et al., 2006, Niki, 2009).

For siRNA or plasmid delivery, $4 \times 10^6$ OSCs were electroporated using one of the following systems: Amaxa 2B (Kit V, program T-029), Amaxa 4D (Buffer SF, program DG150), or MaxCyte (R-50×8 assemblies, program "opt5"). Electroporation mixes contained either 4 µL siRNA (50 µM stock) or 4 µg total plasmid DNA. For MaxCyte electroporations, cells were resuspended in a 1:1 mixture of electroporation buffer and M3 medium. For siRNA transfections a second transfection was performed 48 h after the first, and cells were harvested 96 h after the initial transfection for downstream analyses.

#### Landing site generation and integration

To generate a heterozygous genomic landing site using CRISPR–Cas9–mediated homology-directed repair, candidate sgRNA target sites were computationally identified by intersecting heterozygous single-nucleotide polymorphisms in OSCs with PAM motifs in the dm6 genome, restricting selection to sgRNAs in which the PAM was disrupted on one allele in OSCs to prevent homozygous integration. Off-target potential was assessed using ccTOP with default parameters against the dm6 genome, and a guide targeting locus 86E8 (dm6: chr3R:11529379; OSC: 3R_RagTag:13313192; GGGGGTGGTCGCCCCGATGG) was selected for landing site integration.

The sgRNA was produced by PCR amplification of a long template oligo containing the T7-promoter, the sgRNA targeting sequence and the sgRNA backbone. After gel-purification, the sgRNA was directly loaded into Cas9 protein. A repair template containing the integration cassette (attP–*tj* enhancer–mCherry–P2A–puromycin–*armitage* 3′ UTR–attP) flanked by 500-nt homology arms was generated by PCR and Gibson assembly. Linear donor templates were amplified and purified prior to transfection. Cas9–sgRNA ribonucleoprotein complexes were assembled by pre-incubating purified sgRNA with recombinant Cas9 protein in loading buffer (20 mM HEPES pH 7.5, 150 mM KCl, 0.5 mM DTT, 0.1 mM EDTA) immediately prior to transfection. OSCs were electroporated with Cas9–sgRNA ribonucleoprotein complexes together with 500 ng of linear repair template per transfection, including Alt-R™ Cas9 Electroporation Enhancer and Alt-R HDR Enhancer (IDT), using the Amaxa 4D system. Following recovery, cells were replated and puromycin selection (1:1500) was initiated 2 days post-transfection. After 6 days, Puromycin was removed and single-cells were isolated by fluorescence-activated cell sorting onto irradiated OSC feeder layers (100k OSC per 96 well irradiated with 2x6 Gy with mixing in between, plated 24h prior) the day after. Growing clones were further expanded and maintained under puromycin selection.

Clones were evaluated for correct genomic integration by Southern blotting using probes specific to the landing site and the puromycin cassette, by locus-spanning PCR distinguishing wild-type and targeted alleles, and by digital droplet PCR (ddPCR) using primer sets targeting the cassette and endogenous reference loci.

**Generation of piRNA biogenesis reporters**

All reporter constructs were cloned into an attB integration vector (attB–*tj* enhancer–GFP–experimental 3′ UTR–SV40 UTR–medium enhancer–blasticidin–SV40 3′ UTR–attB) using restriction digestion and Gibson assembly. Constructs were integrated into landing site–containing OSCs by co-transfecting 1 µg of a φC31 integrase expression plasmid together with 3 µg of the attB integration plasmid using Amaxa or MaxCyte electroporation. Following recovery, cells were replated and blasticidin selection (1:250) was initiated 2 days post-transfection. After 6 days, blasticidin was withdrawn and single cells were isolated by fluorescence-activated cell sorting onto irradiated OSC feeder layers. Growing clones were expanded and maintained under blasticidin selection until homogeneous reporter expression was achieved. Successful integration at the landing site was assessed by flow cytometry based on loss of mCherry expression and acquisition of GFP expression.

**Knockdowns in flies for PIWI IPs**

Flies were maintained at 25 °C. Germline knockdown was achieved by crossing UAS-shRNA lines to the maternal triple driver MTD-GAL4 line (#31777; Bloomington Drosophila Stock Center). All shRNA transgenes used in this study were integrated into the attP2 landing site. Zucchini depletion was performed using TRiP.GL00111 (#35227; Bloomington) crossed to MTD-GAL4. As a matched control, an shRNA against *white* (attP2) was crossed to MTD-GAL4.
Adult females were aged for 6 days on apple-juice agar plates supplemented with yeast paste to standardize ovarian morphology prior to dissection.

**Immunofluorescence and Imaging**

To visualize Piwi localization, somatic follicle cells in germline *piwi* knockdown ovaries were stained using a rabbit polyclonal anti-Piwi antibody. Germline Piwi expression was monitored via a germline-specific GFP-Piwi transgene and imaged directly. OSCs were similarly processed for immunofluorescence using the rabbit anti-Piwi antibody. Samples were fixed in 4% paraformaldehyde and imaged on a confocal microscope after antibody staining.

**Small RNA sequencing**

Small RNA sequencing libraries were prepared by Argonaute affinity purification using TraPR resin columns (Grentzinger et al., 2020) or by conventional gel-based small RNA isolation, as indicated for individual experiments (Supplemental Table 2).

*TraPR-based libraries.* For TraPR-based libraries, Argonaute–sRNA complexes were purified as described previously (Grentzinger et al., 2020). OSCs were harvested by trypsinization, washed twice with PBS, snap-frozen in liquid nitrogen, and stored at −70 °C. Cell pellets were lysed in TraPR lysis buffer, clarified lysates were normalized to equal protein input (70–120 µg total protein), and applied to TraPR columns. Argonaute-bound RNAs were eluted, followed by phenol–chloroform extraction and precipitation.

To minimize ligation bias, sequencing adaptors were designed with randomized nucleotides adjacent to the ligating end (Jayaprakash et al., 2011). The 3′ adaptor carried six random nucleotides at its 5′ terminus and internal barcodes (5rApp/NNNNNNXXXX–Adaptor–/3ddC), while the 5′ adaptor carried four random nucleotides at its 3′ terminus. Following 3′ adaptor ligation with T4 RNA ligase 2, truncated KQ (NEB M0373) overnight at 16 °C, samples were pooled as required, spiked with fluorescently labeled, ligation-blocked oligonucleotides as size controls, and gel purified using 12.5% denaturing polyacrylamide–urea gels. RNAs were recovered using the ZR small-RNA PAGE Recovery Kit and subjected to 5′ adaptor ligation using T4 RNA ligase 1.

*Gel-based libraries.* For libraries prepared from total ovarian RNA, 2S rRNA was depleted from 10 µg of input RNA via hybridization to a biotinylated antisense oligonucleotide (5'–Bio-AGTCTTACAACCCTCAACCATATGTAGTCCAAGCAGCACT–3') immobilized on MyOne Streptavidin C1 magnetic beads (Invitrogen). RNA was denatured at 80 °C, hybridized to the bead-bound probe at 50 °C for 1 h, and the rRNA-depleted fraction was recovered by ethanol precipitation. To guide size selection and monitor ligation efficiency, radioactive $\gamma$-$^{32}$P-labeled spike-in oligonucleotides (18 nt and 35 nt) were added to the samples. Small RNAs were isolated by resolving the mixture using a 12.5% polyacrylamide–urea gel and excising the region flanked by the spike-in signals. These spike-ins were ligated to conventional 4N adapters alongside the endogenous small RNA population. Following each ligation step, libraries were gel-purified, with band excision guided by the radioactive signal. After reverse transcription and PCR amplification, the spike-in sequences were removed from the library via restriction digestion. These libraries utilized a non-multiplexed adapter scheme and were pooled only at the final stage using PCR-introduced dual-index barcodes.

*Immunoprecipitation-derived libraries.* Ovaries (600 µl packed volume per genotype) were dissected in ice-cold PBS and lysed in 1× RIPA buffer (50 mM Tris HCl pH 7.5, 150 mM NaCl, 1% Triton X-100, 0.1% SDS, 0.1% sodium deoxycholate, 1 mM EDTA, 0.1 mM Pefabloc). Homogenates were cleared by centrifugation and diluted with 3 ml IP dilution buffer (50 mM Tris HCl pH 7.5, 150 mM NaCl). Antibodies were coupled to M280 sheep anti-mouse IgG Dynabeads (Life Technologies). For Piwi and Aub immunoprecipitations, 150 µl antibody-coupled beads were incubated with 1.5 ml lysate; for Ago3, 300 µl beads were incubated with 3 ml lysate. Lysates were rotated at 4 °C overnight. Beads were collected and washed seven times with IP wash buffer (50 mM Tris HCl pH 7.5, 500 mM NaCl, 2 mM $MgCl_2$, 10% glycerol, 1% Empigen); for Piwi IPs,

150 mM NaCl was used. Bound RNA was isolated by acid phenol–chloroform extraction and ethanol precipitation. For visualization during cloning, 10% of the extracted RNA was labeled with [γ-$^{32}$P]-ATP. The recovered IP-derived small RNAs were used directly as input for the gel-based library preparation pipeline described above.

*Library amplification and sequencing.* For all libraries, reverse transcription was performed after adaptor ligation, followed by PCR amplification with KAPA HiFi polymerase and dual-index barcoded primers. Amplification was monitored in real time using EvaGreen and terminated at 3,500–4,000 RFU to avoid overamplification. Final libraries were size selected on low-melt agarose gels, gel purified (Zymo), quantified by Qubit, pooled, and sequenced (SE50/75/100, Illumina NovaSeq, Hiseq or NextSeq).

**RNA sequencing**

Poly(A)-enriched RNA-seq was performed by extracting total RNA using TRIzol (Invitrogen) from OSCs following siRNA-mediated knockdown. The extracted RNA was treated with DNase I (ThermoFisher Scientific) and purified using the RCC25 kit (Zymo Research) with on-column Zymo DNase I treatment according to the manufacturer's instructions. To ensure high purity of the messenger RNA fraction, the eluted RNA underwent two rounds of poly(A) selection using magnetic Dynabeads Oligo (dT)25 (ThermoFisher Scientific). These samples were then subjected to strand-specific RNA-seq library preparation using the NEBNext Ultra II kit (NEB). Quality control and quantification of both the initial RNA and the final sequencing libraries were evaluated using an Agilent Fragment Analyzer and a Qubit fluorometer (ThermoFisher Scientific).

**TT-SLAM-seq for measuring transcriptional output**

To measure nascent transcription, cells were pulse-labeled for 10 minutes with 4mM 4-thiouridine (4sU) and processed using TT-SLAM-seq (Duffy et al., 2015). Total RNA was extracted using Trizol with chloroform partitioning, precipitated with isopropanol in the presence of DTT and glycogen, and resuspended in nuclease-free water. RNA quality was assessed by Fragment Analyzer, and aliquots were reserved for input (1 µg) and pulldown (50 µg) fractions.

For pulldown samples, total RNA was fragmented to 200–600 nt by heating at 94 °C for 60 s in fragmentation buffer, followed by precipitation with sodium acetate and ethanol. 4sU-labeled transcripts were selectively biotinylated using MTSEA-biotin-XX (MTS-biotin) in a freshly prepared 10× biotinylation buffer (20 mM HEPES pH 7.8, 1 mM EDTA) for 2 h at 25 °C with rotation, protected from light. Unreacted biotin was removed by acid phenol–chloroform extraction, and biotinylated RNA was recovered by ethanol precipitation.

Biotinylated transcripts were enriched using µMACS streptavidin magnetic beads (Duffy et al., 2015). Denatured RNA (65°C, 10 min, snap cool on ice) was incubated with streptavidin beads for 15 min at room temperature, loaded onto pre-equilibrated µMACS columns, washed three times with high-salt buffer (100 mM Tris-HCl pH 7.5, 10 mM EDTA, 1 M NaCl, 0.1% Tween-

20), and eluted twice with 100 mM DTT. Eluted RNA was purified using RNeasy MinElute columns and concentrated to 15 µL.

To introduce 4sU-dependent mismatch signatures for SLAM-seq identification, both input and pulldown fractions were subjected to iodoacetamide (IAA) alkylation. RNA was incubated with 10 mM IAA in 50 mM sodium phosphate buffer (pH 8.0) and 50% DMSO at 50 °C for 15 min; the reaction was then immediately quenched with DTT and the RNA was ethanol precipitated. Following TurboDNase treatment and column purification to remove residual DNA, both fractions underwent rRNA depletion using *Drosophila*-specific oligos and RNase H, as previously described (ElMaghraby et al., 2019). For pulldown samples, the depletion was performed using 1/10th of the oligos and 1/5th of the enzyme concentration used for input samples. The depletion was followed by a final TurboDNase treatment and column-based cleanup.

Strand-specific Illumina sequencing libraries were prepared from rRNA-depleted RNA using the NEBNext Ultra II Directional RNA Library Prep Kit. Input samples were fragmented for 15 min during library preparation, whereas pulldown samples were fragmented for 8 min to account for prior fragmentation and maintain optimal insert sizes (200–600 nt). Libraries were amplified and sequenced on an Illumina platform.

**Generation of anti-Yb monoclonal antibody**

Mouse monoclonal antibodies against Fs(1)Yb (Yb) were generated using a recombinant purified His-tagged Yb antigen (aa 1–200). Hybridoma clones were generated at the MPL Monoclonal Antibody Facility. Hybridoma clones were screened for specificity and high-affinity binding to endogenous Yb in ovarian somatic cells (OSCs).

**Yb Cross-linking Immunoprecipitation (Yb-CLIP)**

CLIP-seq libraries were prepared from semi-confluent OSCs in 15 cm dishes using a protocol modified from Moore et al. (Moore et al., 2014). Cells were washed with PBS and UV-crosslinked on ice (254 nm, 200 mJ/cm$^2$ twice at 90° rotation) using a UV Stratalinker. Harvested cell pellets were lysed in lysis buffer (30 mM HEPES-KOH pH 7.4, 150 mM NaCl, 2 mM $MgCl_2$, 0.5% Triton X-100, 0.2 mM DTT, 0.2 mM Pefabloc) and clarified by centrifugation. Lysates were treated with RNase I (0.08 U per 100 µl cell pellet; Thermo Fisher) for 30 min at room temperature, followed by inhibition with SUPERase In (Thermo Fisher). Immunoprecipitation was performed using monoclonal anti-Yb antibodies coupled to Dynabeads M-280 sheep anti-mouse in TBS-Tx. Beads were incubated with lysates for 4 h at 4 °C and washed four times with high-salt wash buffer (30 mM HEPES-KOH pH 7.4, 1 M NaCl, 2 mM $MgCl_2$, 1% Empigen).

On-bead RNA end repair was performed using T4 polynucleotide kinase (T4 PNK) without ATP to open 2′,3′-cyclic phosphates, followed by 5′-OH radiolabeling with [γ-$^{32}$P]-ATP and final phosphorylation with 10 mM cold ATP. Protein–RNA complexes were eluted in SDS buffer, resolved by 8% SDS-PAGE, and transferred to a 0.45 µm nitrocellulose membrane. Complexes were visualized by autoradiography, and corresponding membrane regions were excised and digested with proteinase K (1 mg/ml). RNA was extracted via acid phenol–chloroform, precipitated, and used for small RNA library preparation. Libraries were generated following standard gel based small RNA cloning and sequenced on an Illumina platform (single-end 50 bp).

**Library Preparation and Sequencing**

Small RNA libraries for CLIP and Input samples were prepared using a low-bias cloning protocol. RNA fragments were ligated to 3′ and 5′ adapters, followed by reverse transcription and PCR amplification. CLIP-seq libraries were sequenced on an Illumina platform. Yb-binding sites were identified by analyzing cross-link-induced mutation signatures (CIMS), specifically focusing on insertion and deletion signatures at cross-link sites to achieve nucleotide-resolution mapping of Yb-RNA interactions. CLIP signals were normalized to the corresponding input and steady-state RNA-seq levels to determine enrichment.

**Computational analyses**

All computational analyses were performed using the *Drosophila* OSC genome (OSC_r1.01) as reference. Bioinformatics analyses utilized standard tools for read alignment, quality control, and feature quantification obtained using the CLIP cluster (https://clip.science). Statistical analyses and modeling were performed in R (v4.3.2). Machine learning models were developed using H2O.ai (v3.46.0.7). Data visualization was performed using ggplot2 and associated packages. A comprehensive list of all software tools with version numbers and citations is provided in Supplementary Table S1.

All computational analyses were performed using the Drosophila OSC genome (OSC_r1.01) as reference. Sequence data processing utilized awk/mawk and standard bioinformatics tools for read alignment (Bowtie v1.3.1, STAR v2.7.10a, minimap2 v2.28-r1221-dirty), quality control, and feature quantification. Statistical analyses and modeling were performed in R (v4.3.2) using tidyverse for data manipulation and mgcv, ppcor, and scam for statistical modeling. Machine learning models were developed using H2O.ai (v3.46.0.7). Data visualization was performed using ggplot2 with extensions for specialized plots and colorblind-accessible palettes. A comprehensive list of all software tools with version numbers and citations is provided in Supplementary Table 1.

<u>Raw-read preprocessing</u>

Illumina reads were processed by removing adapters using cutadapt and filtered for low-complexity sequences using bbduk (entropy = 0.35, entropy window = 18, k = 4). For small RNA sequencing, reads were trimmed of 4 random nucleotides from both ends and filtered for length (>18 nt and ≤35 nt). Reads were first aligned to *Drosophila* rRNA precursor sequences and the mitochondrial genome using Bowtie. Remaining unmapped reads were mapped to the OSC_r1.01 genome using Bowtie (small RNAseq, 1 mismatch) or STAR (RNAseq/TTseq, 3 mismatches). Genomic origin classes were assigned based on overlap with genome annotations, with multi-mapping reads assigned to the annotation class supported by the largest number of alignments.

Reads and alignments were collapsed based on complete sequence identity to facilitate faster processing. For each collapsed sequence, the read name tag encoded sequence identity and abundance, mapping status, annotation assignments (before and after collapsing), and filter status across RNA classes (rRNA, tRNA, mitochondrial, miRNA, pre-miRNA, snRNA, snoRNA, and other ncRNAs).

Small RNA analysis of GFP piRNA biogenesis reporters

Small RNAs mapping to GFP reporter were quantified from small RNA sequencing libraries using a custom computational pipeline (available at [GitHub repository link]). Annotated, collapsed small RNA reads were mapped to sensor plasmid sequences using Bowtie (v1.x) with up to one mismatch allowed, retaining only uniquely mapping reads. Sensor boundaries were defined by aligning reference features (5′ UTR, GFP, experimental 3′ UTR, and SV40 terminator) to each plasmid sequence. For quantification, the sensor region was extracted as the interval spanning 100 nt upstream of the 5′ UTR and 100 nt downstream of the SV40 terminator, with reads mapping to the experimental 3′ UTR (defined as the region between the GFP stop codon and the SV40 element) quantified separately.

Read counts were normalized using one of two strategies depending on the experiment. For experiments with matched qPCR measurements, a qPCR-derived normalization factor was applied to account for differences in sensor expression across samples. For experiments without matched qPCR measurements, counts were normalized to 1 million miRNAs using the total miRNA read count per library.

Per-base coverage tracks were generated using bedtools genomecov for sense and antisense strands separately. Nucleotide composition (A, C, G, T) was computed in 50-nt sliding windows across each sensor using seqkit and faCount. Uridine content within the experimental 3′ UTR was quantified by counting U, UU, and UUU motifs. Plots were generated in R using tidyverse, ggplot2, and cowplot.

Gene and Cluster Region Preparation for downstream analyses

Transcript expression was quantified using StringTie with a merged read set of Illumina RNA-seq and Oxford Nanopore direct RNA sequencing data mapped to annotated gene models. RNA-seq libraries were mapped using STAR with default splice-aware alignment parameters. ONT reads were aligned using minimap2. Transcripts were retained for analysis if they exceeded a TPM threshold of 1 and represented ≥2% of total gene-level TPM. Gene-level expression required a minimum TPM of 2.

Transcript filtering was performed to identify valid coding sequences. Transcript isoforms containing upstream in-frame stop codons (read-through isoforms) were excluded by extracting annotated CDS sequences and identifying transcripts containing stop codons at positions other than the terminal codon. Genes with invalid CDS coordinates or strand-specific overlaps with other expressed genes were excluded. To simplify 3′ UTR isoform analysis, genes were further filtered to retain only those with a dominant stop codon position. For each gene, stop codon positions were determined from all expressed transcript isoforms, and genes were retained only if ≥90% of transcript-level TPM utilized the same stop codon position.

Coding sequence (CDS) blocks were defined as exonic intervals shared across all expressed transcript isoforms of a gene, excluding variable regions specific to individual isoforms. CDS blocks were required to be ≥100 nt in length and ≥75% unique (based on 25-nt k-mer mappability allowing 1 mismatch). CDS blocks overlapping other expressed CDS regions on the same strand were excluded. piRNA cluster regions were divided into 500-nt non-overlapping tiles and required ≥50% uniqueness.

For 3′ UTR isoform determination, UTR boundaries were defined by the dominant stop codon position and the transcript end site(s) identified from ONT direct RNA-seq coverage. Valid 3′ UTR ends were determined by intersecting with ONT reads and filtering low-abundance 3′ ends. Multiple 3′ UTR isoforms were retained if they differed by >200 nt and each represented >5% of total UTR expression. UTRs were filtered to remove those with insufficient representation in major isoforms, those containing in-frame stop codons, and those overlapping transposable elements or other expressed genes.

CDS blocks and piRNA cluster tiles were quantified using strand-specific read coverage from RNA-seq and TT-seq (transient transcriptome sequencing) libraries. Read counts were normalized to reads per kilobase per million mapped reads (RPKM). CDS blocks were required to have ≥0.1 RPKM in both RNA-seq and TT-seq to be retained for analysis. Gene-level expression was determined exclusively from CDS block RPKM values.

For 3′ UTR isoform quantification, expression levels were not measured directly from UTR coverage. Instead, 3′ UTR isoform abundance was derived by scaling the CDS block expression level according to the relative abundance of transcript 3′ ends identified from Nanopore direct RNA-seq.

Final output tables were generated to facilitate downstream analyses. CDS block tables included genomic coordinates, uniqueness annotations, expression levels (RPKM) from RNA-seq and TT-seq, and filter tags indicating excluded regions (overlapping genes, intron-spanning, transposable element overlaps). piRNA cluster tile tables contained genomic coordinates, uniqueness information, and strand-specific expression values. 3′ UTR isoform tables included UTR boundaries, dominant stop codon positions, relative isoform abundances, and filter annotations. All tables were merged with corresponding headers and exported in tab-delimited format for integration with downstream analyses. Visualization tracks were generated in UCSC-compatible formats (BigBed and BigWig) for RNA-seq and TT-seq RPKM values.

<u>Creation of replicate-averaged libraries (small RNA-seq, RNA-seq, and TT-seq)</u>

Before the creation of replicate-average libraries, RNA-seq and TT-seq reads were re-aligned to the genome. Raw reads were trimmed of 6 bases from both the 5′ and 3′ ends using seqkit. Trimmed reads were aligned to the *Drosophila* genome (dm6) using STAR (version 2.7.10a) in two-pass mode with local alignment enabled. Post-alignment, STAR alignments were annotated as unique or multi-mapping reads within the name-tag.

Replicate libraries for RNA-seq, TTseq and small RNA-seq datasets were merged into genotype-level "average" libraries through read normalization and averaging. The original read count embedded in the read name was parsed and converted into a scaled per-read score equal to count / NORM / nLIB, ensuring that each replicate contributes equally to the final averaged signal. For RNA-seq and TT-seq, reads were normalized to their respective genome-uniquely aligning read counts (reads per 10 million reads RPM). For small RNA-seq, reads were normalized to miRNA counts (reads per 1 million miRNA reads $RPM_{miR}$). These normalized, per-replicate BED files were then concatenated and sorted to generate a single replicate-averaged BED representation for each genotype.

For small RNA-seq, additional steps were included to recover informative piRNA reads originating from repetitive piRNA cluster regions. Because piRNAs are generally assumed to arise predominantly from annotated piRNA clusters, many reads mapping to multiple genomic locations can still be assigned unambiguously within these cluster intervals. To retain such reads, a "cluster-unique" set was computed by intersecting all multi-mapper-averaged reads with piRNA cluster annotations and selecting those mapping only once within the combined cluster intervals, regardless of total genome-wide alignments. The resulting cluster-unique reads were combined with conventional genome-unique reads and merged into a single set.

CLIP-seq preprocessing and crosslink-site extraction

CLIP-seq replicates were merged using samtools merge. Reads were partitioned into indel-containing versus non-indel alignments based on the CIGAR string (presence of I or D). For each subset (all reads and indel reads), alignments were converted to BED format, and read counts were recovered from the read-name annotation and stored in the 5th column of the BED file. Uniquely mapped reads (mapping=u) were written as both full-length and 3′-end–collapsed BED files.

To characterize crosslinking sites, genomic "xLink sites" were derived for each alignment. For indel-containing reads, deletion positions were parsed from the CIGAR string to approximate the crosslink site; for all reads (and separately for no-indel reads), the site was approximated by the alignment midpoint. Sites were expanded by ±10 nt, converted to BED format, and intersected with annotated 3′ UTR intervals. Corresponding UTR-centered sequences were extracted from the genome using bedtools getfasta. Motif enrichment analysis was performed using STREME (MEME Suite) in RNA mode with motif widths ranging from 3 to 5 nt, using CLIP input sequences as the negative control set.

Isoform-abundance scaling of small RNA libraries

For analyses requiring 3′ UTR-resolved signals, small RNA libraries were further processed to correct for differences in 3′ UTR isoform abundance. Because longer transcript isoforms are typically less abundant than the most upstream isoform, small RNA counts mapping to downstream UTR regions were scaled upward accordingly. This adjustment ensures that small RNA signals can be compared uniformly across the entire 3′ UTR, independent of isoform-specific expression differences.

Size-profile, 1U enrichment, and phasing analysis

Size-profile, 1U-enrichment, and phasing analyses were performed by selecting small RNA reads according to genotype- and annotation-specific categories as defined in the figure panels. For size profiles, normalized read counts were summed by observed piRNA length and visualized either as absolute normalized counts or as fraction per-length distributions restricted to the 18–35 nt window.

For 1U-enrichment analysis, all ≥23 nt non-miRNA reads were uncollapsed and converted to FASTA format. Sequence logos were generated using WebLogo (Crooks et al., 2004) from abundance-weighted FASTA files (each sequence expanded proportional to its read count) to assess position-specific nucleotide preferences and detect 1U biases.

For phasing analysis, genome-wide 5′ and 3′ ends of uniquely mapping piRNAs were extracted and summarized at 1 nt resolution. Reads sharing identical genomic coordinates were aggregated to obtain per-locus end-counts. The 3′ ends of abundant piRNAs (>10 counts) were intersected with nearby 5′ ends on the same strand within a ±10 nt window to compute the distribution of observed 3′→5′ offsets. These distances were compiled into positional histograms representing the relative frequency of offsets from −10 to +10 nt. A phasing score (Z score) was calculated as the standardized deviation at +1 nt relative to the local background distribution.

Genomic source annotation and quantification of RNA-reads

Small RNA reads ≥23 nt or RNA-seq reads, excluding reads annotated as miRNAs, rRNAs, tRNAs, snoRNAs, snRNAs, mitochondrial RNAs, ncRNAs, and pseudogenes, were extracted from collapsed 3′-end BED files for each library. Filtered reads were intersected with a reference annotation BED file using bedtools intersect (-wao -s -sorted) to assign genomic source annotations to each read based on strand-specific overlap.

For reads overlapping multiple annotation features, a priority-based assignment rule was applied. Reads were grouped by unique sequence identifier, and the annotation with the highest number of overlapping instances was selected. In cases of tied overlap counts, priority was determined by a predefined ruleset. Reads were additionally flagged if they overlapped with defined piRNA cluster regions.

Read counts were aggregated by annotation category (5′ UTR, CDS, 3′ UTR, intron, transposable element, antisense transposable element, or none) and cluster status (Y/N). The resulting quantification table was used to generate stacked bar charts and alluvial plots showing the proportional distribution of small RNA reads across genomic features between libraries as indicated in the figure panels.

Full-length UTR/CDS analysis

For quantification of small RNA signal across complete transcript regions, uniquely mapped, isoform-corrected 3′-end reads were intersected with a reference set of CDS blocks and 3′ UTRs. From each library, reads were restricted to uniquely mapping small RNAs ≥23 nt and filtered to exclude reads annotated as rRNA precursor, tRNA, snoRNA, snRNA, mitochondrial RNAs, or miRNAs. Using strand-specific overlap, small RNA coverage was quantified for each annotated transcript interval.

For each transcript (FBgn), several summary metrics were calculated from the full-length coverage profile using length and mappability information from the CDS/UTR annotation table:

- the total normalized small RNA signal across the full interval,
- signal normalized by the transcript's uniquely mappable fraction (UNIQ) and length, and
- additional summary measures for regions beyond defined distances from the CDS start (e.g. >250 nt, >400 nt, >500 nt into the interval), including UNIQ-normalized signal and the median per-position coverage within the uniquely mappable fraction.

To obtain RNA-abundance–corrected estimates of small RNA targeting, individual read values were divided by the RNA-seq RPKM of the corresponding transcript before output. Genomic coordinates of 3′-end reads were then adjusted to reflect the true end position of each small RNA (extending coordinates by the inferred read length upstream or downstream, depending on strand). The resulting RNA-seq–corrected 3′-end BED files were sorted and converted into strand-specific bedGraph and bigWig tracks for visualization and downstream analyses.

Downstream analyses of full-length CDS and 3′ UTR piRNA production, RNA/TT-normalization ("RNA-norm" and "TT-norm") and biogenesis-efficiency comparisons between transcript classes (CDS, UTR, cluster-derived) were performed using these quantified transcript-level tables together with the RNA-seq and TT-seq expression estimates described above. For biogenesis-factor perturbation experiments, small RNA counts from each knockdown condition were compared to the corresponding siGFP control on a per-transcript basis (e.g. fold-changes in CDS-, UTR-, and cluster-derived intervals, and global losses of total and cluster-derived piRNA output). Relationships between biogenesis efficiency and Yb dependency were then evaluated by restricting to transcripts with detectable small RNA signal in both wild type and Yb depleted libraries. Yb dependency was defined as the ratio of wild type to Yb depleted small RNA counts, biogenesis efficiency as wild type small RNA counts normalized by RNA seq RPKM, and correlations were assessed using Pearson partial correlations (controlling for wild type small RNA levels), linear regression, and Kendall's $\tau$.

<u>Parameter-sweep</u>

To systematically optimize the parameters used to quantify piRNA biogenesis efficiency and Yb dependency along mRNA 3′ UTRs, we implemented a parameter-sweep framework. Genomic intervals corresponding to 3′ UTRs were partitioned into fixed-length, non-overlapping 100-nt tiles. For each tile, small RNA reads reads were quantified and a piRNA biogenesis efficiency score was computed as the ratio of small RNA density (from the 100-nt tile) to RNA-seq RPKM. To evaluate the impact of local sequence features upstream and downstream of the piRNA quantification tile, we defined a series of "nucleotide-evaluation" windows by extending each tile (50–800 nt, in 50-nt increments) and systematically shifting these windows relative to the piRNA tile (−800 to +100 nt, in 50-nt increments). For each nucleotide-evaluation window, mono- and dinucleotide occurrences were quantified. Windows extending beyond UTR boundaries were either evaluated using the adjacent CDS sequence or excluded from the analysis. Correlation analyses between piRNA biogenesis efficiency and nucleotide features were restricted to tiles that remained analyzable across all combinations of window shifts and extensions. This common set of shared tiles was further filtered based on minimum coverage, sequence uniqueness, and length. For the resulting high-confidence tile set, correlations between biogenesis efficiency and individual nucleotide features were computed in R and aggregated into a merged summary table. Correlation heatmap plots were generated using the tidyverse, cowplot, and ggplot2 packages in R.

Tile analysis

To quantify piRNA biogenesis efficiency and Yb dependency along CDS, 3′ UTRs and piRNA clusters, we generated 100-nt non-overlapping genomic tiles spanning each region. Each tile was annotated with genomic coordinates, transcript identity, length, sequence uniqueness (mappability) and RNA-seq and TT-seq coverage derived from the scaled CDS-block quantification (RPKM). For tiles overlapping piRNA clusters, a cluster-specific uniqueness metric was computed to account for reads that map to multiple genomic locations but are unique within the piRNA cluster itself. Quantification was performed separately for genome uniquely mapping reads and for the cluster-unique extended read sets.

Small RNA-seq and CLIP libraries (both 3′ UTR isoform corrected) were quantified for each 100-nt tile. For each library, small RNA reads ≥23 nt (or all reads for CLIP datasets) were filtered to exclude rRNA, tRNA, snoRNA, snRNA, mitochondrial RNA, and miRNA annotations. Filtered reads were intersected with tile coordinates using bedtools intersect (strand-specific mode, -s; split mode for spliced alignments, -split), and the total number of overlapping nucleotides was summed per tile. Isoform corrected read counts were length normalized by tile length and sequence uniqueness to yield a uniqueness-corrected read density (reads per 100 uniquely mappable nucleotides).

To evaluate the impact of sequence features on piRNA biogenesis, nucleotide content (mono- and dinucleotide frequencies) was calculated in two complementary ways: (i) within the 100-nt piRNA quantification tiles themselves (Fig. 2/5), and (ii) within extended nucleotide-evaluation tiles (750 nt, shifted −600 nt relative to each 100-nt piRNA tile) as determined in the parameter-sweep (Fig. 3/4).

For each tile, biogenesis efficiency was computed as the ratio of small RNA density (from the 100-nt tile) to RNA-seq RPKM (RNA-normalized) or TT-seq RPKM (transcription-normalized). Quality filters required RNA-seq coverage ≥5 RPKM and sequence uniqueness >50%.

Motif discovery based on tile sequences

For motif discovery, nucleotide sequences corresponding to the 100-nt evaluation windows of high- and low-efficiency tiles were extracted from the genome using seqkit. To increase motif resolution, each sequence was fragmented into overlapping 10-nt windows with a 5-nt step using seqkit sliding. Discriminative motif discovery was performed using STREME (MEME Suite), comparing high-efficiency sequences against low-efficiency sequences as background. Motif searches were conducted for motifs ranging from 3 to 5 nt (--minw 3 --maxw 5) in RNA mode (--rna).

<u>Machine learning model of piRNA biogenesis efficiency in wt or Yb depleted conditions</u>

To identify the sequence and expression features that determine piRNA biogenesis efficiency, we trained gradient boosting machine (GBM) models using the H2O AutoML framework in R. The analysis was performed on 100-nt tiles from 3′ UTRs with associated 750-nt nucleotide-evaluation windows (shifted −600 nt). Input features included dinucleotide frequencies within the extended windows, RNA-seq and TT-seq expression levels (RPKM), UTR length, and tile position within the UTR. The target variable was $\log_{10}$-transformed small RNA density in wild-type or Yb-depleted samples. Tiles were filtered to require RNA-seq ≥1 RPKM, TT-seq ≥1 RPKM, and sequence uniqueness >50%.

To prevent data leakage and ensure generalizability, tiles were split into training (80%), validation (10%), and test (10%) sets at the transcript level, such that all tiles from a given 3′ UTR were assigned to the same partition. To address the imbalanced distribution of piRNA production levels, training samples were weighted using an exponential function of small RNA density: weight = 100^(1/((max_sRNA + 1)/(tile_sRNA + 1))), which upweighted tiles with higher piRNA production. Models were trained using H2O AutoML with up to 30 GBM candidates, optimized for root mean squared error (RMSE) on the validation set, and evaluated on the held-out test set using Pearson correlation ($R^2$), concordance correlation coefficient (CCC), and Spearman's ρ.

To assess feature importance, we performed an ablation study by training six model variants: (i) all features, (ii) without dinucleotide content, (iii) without RNA-seq, (iv) without TT-seq, (v) without RNA-seq and TT-seq, and (vi) without positional information. Variable importance was quantified using scaled importance scores from the best-performing model, and SHAP (SHapley Additive exPlanations) values were computed to interpret individual feature contributions.

To evaluate model generalizability, we applied the full-feature model trained on 3′ UTR tiles to predict piRNA production from CDS tiles (test set only, matched by transcript). Because CDS regions exhibited systematically lower piRNA production than UTRs, we computed a calibration offset as the mean difference between observed and predicted $\log_{10}$ small RNA density in CDS tiles and applied this correction to CDS predictions. Model performance on CDS tiles was assessed using the same metrics as for UTR tiles.

<u>Positional analysis of piRNA biogenesis along 3′ UTRs</u>

To assess whether piRNA biogenesis efficiency varies with position along 3′ UTRs, we analyzed the relationship between tile position and small RNA production. We selected 3′ UTR tiles meeting quality filters (RNA-seq ≥5 RPKM, TT-seq ≥5 RPKM, sequence uniqueness >50%, UTR length >700 nt, and tile position nPOS >0). For each tile, transcription-normalized small RNA density was calculated as the ratio of small RNA counts to TT-seq RPKM. To compare piRNA production dynamics across UTRs of different lengths and expression levels, we computed a scaled piRNA level for each tile by normalizing to the maximum small RNA density within the same UTR.

Positional trends were visualized by plotting transcription-normalized small RNA density as a function of tile position (tiles 1–40 from the 5′ end of the 3′ UTR). For each position, the distribution of piRNA production across all UTRs was summarized using boxplots, and a smoothed trend line was fitted using locally weighted regression (LOESS). The number of tiles contributing to each position was annotated on the plot. This analysis revealed the characteristic "ramp-up" pattern of piRNA biogenesis along 3′ UTRs, with piRNA production increasing progressively from the 5′ to 3′ end of the UTR.

Application of the UTR-trained model to piRNA clusters

To test whether the sequence and expression features that determine piRNA biogenesis in 3′ UTRs also apply to piRNA clusters, we applied the UTR-trained GBM model to predict piRNA production from tiles in the *flamenco* piRNA cluster. Because piRNA clusters lack the stable RNA-seq and TT-seq signals characteristic of protein-coding transcripts, we first modeled the position-dependent decay of transcription along *flamenco*. For tiles with sequence uniqueness ≥50% and detectable TT-seq signal (>1 RPKM), we fit a monotonic decreasing smooth curve to $\log_{10}$-transformed TT-seq RPKM as a function of genomic position using shape-constrained additive models (SCAM) with a monotonic P-spline basis (*bs* = "mpd", smoothing parameter = 0.1). A similar model was fit to RNA-seq RPKM for tiles beyond position 2,500 nt. Predicted TT-seq and RNA-seq values were then used as input features for tiles where measured expression was unreliable or absent.

We applied three model variants to *flamenco* tiles (positions 5,000–600,000 nt; tiles located within the first 5,000 nt were excluded due to the presence of a strongly spliced intron; cluster-unique mappability >50%): (i) the full-feature model, (ii) the model without dinucleotide content, and (iii) the model without TT-seq. Because piRNA clusters exhibited systematically different piRNA production levels than UTRs, we computed a linear calibration offset by regressing observed $\log_{10}$ small RNA density against predicted values and applied this correction to all cluster predictions. Model performance was evaluated using $R^2$, CCC, Pearson's $r$, and Spearman's $\rho$. To visualize spatial patterns of piRNA production along *flamenco*, we computed a rolling average of observed and predicted small RNA density using a 20-tile sliding window. Gaps >5,000 nt were treated as discontinuous regions. Observed and predicted piRNA levels were plotted as a function of genomic position, with smoothed trend lines overlaid to highlight regional variation in piRNA biogenesis efficiency.

Germline piRNA analysis

To identify germline-specific piRNA-producing transcripts, we selected genes not expressed in somatic OSC cells. From the full transcriptome annotation, we identified genes with a single stop codon and a single 3′ UTR end by grouping transcripts by gene ID and filtering for unique stop codon positions (for plus-strand genes: CDS end; for minus-strand genes: CDS start) and unique

3′ UTR termini. For each gene, we extracted the longest CDS and 3′ UTR isoforms and combined these with piRNA cluster regions to generate a unified set of germline-expressed regions. Sequence uniqueness was calculated for each region by intersecting with 25-mer uniqueness tracks (allowing 1 mismatch for small RNA mapping) using bedtools intersect, and the percentage of uniquely mappable sequence was computed.

Small RNA-seq and RNA-seq libraries were quantified over these germline regions using bedtools intersect. For small RNA-seq, reads were filtered to retain uniquely mapping reads ≥23 nt, excluding rRNA, tRNA, snoRNA, snRNA, mitochondrial, and miRNA annotations. For each region, total read counts were summed across all overlapping tiles, and read density was normalized by sequence uniqueness (reads per kilobase of unique sequence, RPKU). RNA-seq libraries were quantified similarly, but without length filtering. Quantifications were performed in parallel across multiple libraries, and results were merged into a single table containing gene ID, region length, sequence uniqueness, and normalized read counts for all libraries.

To distinguish germline (GL) from soma-enriched piRNA-producing loci, we quantified piRNA abundances from control ovarian somatic cells (sh-*white*) and germline-depleted ovaries (sh-*piwi* PiwiIP). Libraries were normalized by calculating the median ratio between replicate datasets and applying this factor to equalize distributions. For each genomic feature (CDS, 3′ UTR, or piRNA cluster), we calculated the soma-to-germline ratio as: $\text{Ratio}_{\text{soma/GL}} = \text{piRNA}_{\text{Wsh}}/\text{piRNA}_{\text{shPiwi}}$. Features with ratios $> 5$ were classified as germline-enriched, those with ratios $< 0.2$ as soma-enriched, and intermediate values as mixed. Only features with $> 200$nt length and detectable piRNA levels in both conditions were analyzed.

To assess dependency on specific biogenesis factors, we calculated fold-changes in piRNA abundance upon knockdown of *aub/ago3* or *zuc* relative to wild-type controls. For *aub/ago3* dependency, we computed: $\text{Ratio}_{\text{AubAGO3}} = \text{piRNA}_{\text{AubAGO3sh}}/\text{piRNA}_{\text{Wsh}}$. For *zuc* dependency, ratios were calculated separately for Piwi-IP and Aub-IP samples: $\text{Ratio}_{\text{Zuc}} = \text{piRNA}_{\text{shZuc}}/\text{piRNA}_{\text{shWhite}}$. Results were visualized using beeswarm plots.

Germline ping-pong analysis

To analyze piRNA signatures across the transcriptome, an "extended transcriptome" was generated by merging *Drosophila* transcripts (FlyBase r6.66) with TE consensus sequences. To ensure mapping specificity, minimap2 was used to exclude transcripts with $> 100$bp of TE homology or significant gene-gene overlaps ($> 70\%$ sense or any antisense), retaining only the longest isoform per gene. Small RNA reads ($\geq 23$ nt) from Piwi, Aub, and Ago3 IPs were aligned to this reference using bowtie (-v 1 -a), excluding structural RNAs and miRNAs. Alignment coordinates were converted into a nucleotide-resolution matrix of normalized 5′ and 3′ end frequencies for both strands using bedtools and custom awk scripts. The 10-nt 5′–5′ overlap signature was then quantified using a modified Brennecke linkage analysis, calculating the probability of a 10-nt antisense offset within a 20-nt window. The significance of this signature was determined via Z-score: $Z = (P_{10} - \mu)/\sigma$, where $\mu$and $\sigma$represent the mean and standard deviation of all other offsets in the window. These calculations were performed in R using data.table and mclapply, with final results filtered for a responder score and median background $> 0.01$.

Germline transposon analysis

To quantify piRNA production from individual transcripts and TEs, small RNA reads (≥ 23 nt) from wild-type and *aub*/*Ago3* knockdown libraries were aligned to the extended transcriptome using bowtie (-v 1 -a), excluding structural RNAs and miRNAs. Strand-specific read counts were summed for each transcript to generate a quantification matrix containing sense and antisense piRNA abundances. Strand ratios (sense/antisense) were calculated to assess directional biases in piRNA production. To examine spatial distribution of piRNAs along TEs, nucleotide-resolution coverage profiles were generated using custom awk scripts that accumulated strand-specific read counts at every position. Coverage data were merged across libraries and visualized in R using ggplot2, with sense coverage plotted as positive values and antisense as negative. Deregulated TEs (identified from RNA-seq analysis) were compared to unchanged TEs to assess changes in strand ratio and positional coverage patterns upon knockdown of ping-pong factors.

Obtaining Drosophila Gypsy/Gypsy, 17.6/Gypsy and MDG1/Gypsy sequences from RepBase

Sequences of Gypsy LTR retrotransposons from Drosophila species were obtained from RepBase in January 2026. Firstly, we used hmmer 3.3.1 (http://hmmer.org/) to classify annotated open reading frames (ORFs) into subgroups by comparing them against ORFs of known retroelements that are available from the Gypsy Database 2.0 (https://gydb.org/index.php/Main_Page). The Gypsy Database 2.0 collects hmm profiles of Ty1/Copia, Bel/Pao, Ty3/gypsy, and Retrovirus ORFs. We took Gypsy LTR elements with a POL ORF that has both RT and INT domains, most closely related to Ty3/gypsy. Followingly, we made a multiple sequence alignment of the RT domains using mafft-7.505 with the option '--auto' and inferred the phylogenetic relationships using iqtree-1.6.12 with the option '-m rtREV+R4 -bb 1000'. The phylogenetic tree largely agreed with the hmmer results and allowed us to obtain a total of 199 Drosophila LTR elements from Gypsy/Gypsy, 17.6/Gypsy and MDG1/Gypsy groups. The multiple sequence alignment and the phylogenetic tree were made available at (https://github.com/RippeiHayashi/Yb_2026). Next, we used hmmer and hhpred (https://toolkit.tuebingen.mpg.de/tools/hhpred) to classify annotated ORFs from the 199 Gypsy LTR elements and kept 512 ORFs that were predicted to be GAG, POL and ENV ORFs for further analyses. Nucleotide and amino acid sequences of the 512 ORFs that were used to measure codon frequencies and nucleotide compositions were made available at (https://github.com/RippeiHayashi/Yb_2026). The classification of individual LTR elements and the domain prediction of ORFs are summarised in table S3.

Measuring nucleotide compositions of *Drosophila melanogaster* protein coding mRNAs and *Gypsy* LTR retrotransposon sequences in Drosophilids

We obtained coding sequences (CDS) of D. melanogaster protein coding genes from Ensembl release 115 (equivalent of FlyBase r6.54). We obtained 3′ untranslated region (UTR) sequences

using the genomic coordinate information from the gtf file. We took the longest CDS and 3′ UTR sequences per gene, excluding sequences shorter than 600 and 300 nucleotides for CDS and 3′ UTR, respectively. CDS and 3′ UTR sequences were grouped into 50 bins across their length, and the nucleotide compositions were measured in each bin to produce line plots using ggplot2_3.5.2. We trimmed the START and STOP codons from the CDS for this analysis.
We measured nucleotide compositions of the internal region of 199 Drosophila Gypsy/Gypsy, 17.6/Gypsy and MDG1/Gypsy elements obtained above and produced a heatmap using ggplot2. We also measured the nucleotide compositions of aforementioned 512 Gypsy ORFs in a line plot after grouping the sequences into 50 bins across their length.

<u>Measuring codon frequencies and inferring expected adenosine contents of Gypsy ORFs and protein coding mRNAs</u>
We measured codon frequencies using the non-redundant D. melanogaster CDSs (longer than 600 nucleotides) mentioned above, which yielded expected adenosine frequencies per nucleotide per amino acid codons. For example, isoleucine codons are used in the following frequencies: 0.010 (ATA), 0.022 (ATC) and 0.17 (ATT). This translates to an average of 0.40 adenosine per nucleotide per isoleucine codon, slightly more than 0.33 because of the ATA codon. We measured expected adenosine frequencies of aforementioned 512 Gypsy ORFs and D. melanogaster CDSs. We made a contour plot of expected and observed adenosine frequencies per element/gene using ggplot2.

**Table S1. Software Tools used for data analysis**

| Name | Version | Reference | Description of use | Category |
|---|---|---|---|---|
| **Bowtie** | 1.3.1 | (Langmead et al., 2009) | Small RNA and genomic read alignment (allowing up to 1 mismatch) | Bioinformatics - Alignment |
| **minimap2** | 2.28-r1221-dirty | (Li, 2018) | Purpose: Oxford Nanopore direct RNA-seq alignment | Bioinformatics - Alignment |
| **STAR** | 2.7.10a | (Dobin et al., 2013) | RNA-seq alignment | Bioinformatics - Alignment |
| **MEME Suite** | 5.5.7 | (Bailey, 2021) | Discriminative motif discovery using STREME (RNA mode, minw 3, maxw 5) | Bioinformatics - Motif Analysis |
| **Weblogo** | 3.7.12 | (Crooks et al., 2004) | Sequence logo generation for position-specific nucleotide preferences | Bioinformatics - Motif Analysis |
| **bbduk (BBTools)** | 39.13 | (Bushnell, 2014) | Low-complexity filtering (entropy = 0.35, entropy window = 18, k = 4) | Bioinformatics - Preprocessing |

| | | | | |
|---|---|---|---|---|
| **cutadapt** | 4.9 | (Martin, 2011) | 3' adapter removal from sequencing reads | Bioinformatics - Preprocessing |
| **StringTie** | 2.2.0 | (Pertea et al., 2015) | Transcript assembly and quantification | Bioinformatics - RNA-seq |
| **bedtools** | 2.28.0 | (Quinlan and Hall, 2010) | Genomic interval operations, coverage tracks (genomecov), BAM to BED conversion, sequence extraction (getfasta), interval intersections | Bioinformatics - Utilities |
| **samtools** | 1.13 | (Danecek et al., 2021) | BAM file manipulation, merging replicates | Bioinformatics - Utilities |
| **seqkit** | 2.9.0 | (Shen et al., 2024) | Nucleotide composition analysis, sequence manipulation | Bioinformatics - Utilities |
| **faCount (UCSC Genome Browser)** | v479 | (Perez et al., 2025) | Nucleotide counting | Bioinformatics - Utilities |
| **SLURM** | 24.05.3 | (Jette and Wickberg, 2023) | Job scheduling and workload management | Infrastructure - Cluster Computing |
| **Apptainer** | 1.1.9-1 | (Developers, 2021, Kurtzer et al., 2017) | Container management for reproducible computing | Infrastructure - Containerization |
| **mawk** | 1.3.4 20240905 | | Fast AWK implementation for text processing, data filtering, normalization calculations, and formatting | Bioinformatics - Utilities |
| **R** | 4.3.2 | | Statistical analysis, visualization, machine learning | Programming Language |
| **tidyverse** | 2.0.0 | (Wickham et al., 2019) | Data manipulation and analysis framework (includes dplyr, tidyr, readr, etc.) | R Package - Data Manipulation |
| **dplyr** | 1.1.4 | (Wickham et al., 2023a) | Data manipulation (filtering, grouping, summarizing) | R Package - Data Manipulation |
| **forcats** | 1.0.0 | (Wickham, 2023a) | Factor manipulation | R Package - Data Manipulation |
| **readr** | 2.1.5 | (Hadley Wickham, 2023) | Fast reading of rectangular data | R Package - Data Manipulation |
| **stringr** | 1.5.1 | (Wickham, 2023b) | String manipulation | R Package - Data Manipulation |

| | | | | |
|---|---|---|---|---|
| **tibble** | 3.2.1 | (Kirill Müller, 2023) | Modern data frames | R Package - Data Manipulation |
| **tidyr** | 1.3.1 | (Hadley Wickham, 2024) | Data tidying and reshaping | R Package - Data Manipulation |
| **H2O.ai** | 3.46.0.7 | (Tomas Fryda, 2025) | Gradient boosting machine (GBM) models for piRNA biogenesis prediction, AutoML framework | R Package - Machine Learning |
| **Metrics** | 0.1.4 | (Frasco, 2018) | Machine learning model evaluation metrics | R Package - Machine Learning |
| **mgcv** | 1.9.0 | (Wood, 2017) | Generalized additive models (GAMs) | R Package - Statistics |
| **ppcor** | 1.1 | (Kim, 2015) | Partial correlation analysis controlling for confounders | R Package - Statistics |
| **scam** | 1.2-18 | (Pya, 2025) | Shape-constrained additive models | R Package - Statistics |
| **DescTools** | 0.99.59 | (Signorell, 2025) | Descriptive statistics and data manipulation | R Package - Statistics |
| **MASS** | 7.3-60 | (W. N. Venables, 2002) | Statistical functions and datasets | R Package - Statistics |
| **zoo** | 1.8-13 | (Achim Zeileis, 2005) | Time series and irregular data handling | R Package - Statistics |
| **devtools** | 2.4.5 | (Hadley Wickham, 2002) | Package development and installation from GitHub | R Package - Utilities |
| **cowplot** | 1.1.3 | (Wilke, 2024) | Figure composition and publication-ready plot layouts | R Package - Visualization |
| **ggbeeswarm** | 0.7.2 | (Clarke et al., 2023) | Beeswarm plots (categorical scatter plots) | R Package - Visualization |
| **ggforce** | 0.4.2 | (Pedersen, 2024) | Extended ggplot2 functionality (faceting, shapes, etc.) | R Package - Visualization |
| **ggplot2** | 3.5.2 | (Wickham, 2016) | Primary plotting and data visualization | R Package - Visualization |
| **ggpubr** | 0.6.0 | (Kassambara, 2023) | Publication-ready plots, statistical comparisons | R Package - Visualization |
| **ggrastr** | 1.0.2 | (Viktor Petukhov, 2023) | Rasterization of ggplot2 layers for large datasets | R Package - Visualization |
| **plotly** | 4.10.4 | (Sievert, 2020) | Interactive visualizations | R Package - Visualization |
| **ggalluvial** | 0.12.5 | (Jason Cory | Alluvial/Sankey diagrams | R Package - Visualization |

| | | | | |
|---|---|---|---|---|
| | | Brunson, 2023) | | |
| **ggalt** | 0.4.0 | (Bob Rudis, 2017) | Additional coordinate systems and geoms for ggplot2 | R Package - Visualization |
| **gghighlight** | 0.4.1 | (Yutani, 2023) | Highlight specific data points in plots | R Package - Visualization |
| **ggokabeito** | 0.1.0 | (Barrett, 2021) | Okabe-Ito colorblind-safe color palette | R Package - Visualization |
| **khroma** | 1.16.0 | (Frerebeau, 2025) | Color schemes for scientific data visualization | R Package - Visualization |
| **paletteer** | 1.6.0 | (Hvitfeldt, 2021) | Comprehensive collection of color palettes | R Package - Visualization |
| **scales** | 1.3.0 | (Wickham et al., 2023b) | Scale functions for visualization | R Package - Visualization |
| **scico** | 1.5.0 | (Thomas Lin Pedersen, 2023) | Scientific color palettes (perceptually uniform, colorblind-safe) | R Package - Visualization |

**Table S2 (separate file). NGS datasets used for data analysis**

All datasets published or created for this study, with information including sequencing and library preparation details, normalization factors, Figure-panel references.

**Table S3 (separate file). Classification and domain annotation of *Drosophila* Gypsy LTR retrotransposons.**

Containing information including nucleotide compositions of the internal region and the domain prediction of the encoded proteins.
